# PIN2-mediated self-organizing transient auxin flow contributes to auxin maxima at the tip of Arabidopsis cotyledons

**DOI:** 10.1101/2024.06.24.599792

**Authors:** Patricio Pérez-Henríquez, Shingo Nagawa, Zhongchi Liu, Xue Pan, Marta Michniewicz, Wenxin Tang, Carolyn Rasmussen, Jaimie Van Norman, Lucia Strader, Zhenbiao Yang

## Abstract

Directional auxin transport and formation of auxin maxima are critical for embryogenesis, organogenesis, pattern formation, and growth coordination in plants, but the mechanisms underpinning the initiation and establishment of these auxin dynamics are not fully understood. Here we show that a self-initiating and - terminating transient auxin flow along the marginal cells (MCs) contributes to the formation of an auxin maximum at the tip of Arabidopsis cotyledon that globally coordinates the interdigitation of puzzle-shaped pavement cells in the cotyledon epidermis. Prior to the interdigitation, indole butyric acid (IBA) is converted to indole acetic acid (IAA) to induce PIN2 accumulation and polarization in the marginal cells, leading to auxin flow toward and accumulation at the cotyledon tip. When IAA levels at the cotyledon tip reaches a maximum, it activates pavement cell interdigitation as well as the accumulation of the IBA transporter TOB1 in MCs, which sequesters IBA to the vacuole and reduces IBA availability and IAA levels. The reduction of IAA levels results in PIN2 down-regulation and cessation of the auxin flow. Hence, our results elucidate a self-activating and self-terminating transient polar auxin transport system in cotyledons, contributing to the formation of localized auxin maxima that spatiotemporally coordinate pavement cell interdigitation.

## Introduction

Proper formation and morphogenesis of an organ requires not only the differentiation of cells and tissues, but also their coordinated growth in time and space, but the mechanisms for this coordination are mostly unexplored in multicellular organisms. The Arabidopsis cotyledon is an outstanding and trackable example of growth coordination and morphogenetic pattern formation towards the building of an efficient light-capturing flat structure. The expanding epidermal layers of pavement cells (PCs) on both sides of the flat structure accommodate the expanding mesophyll cells. On the edges, a file of elongated cells called the marginal cells (MCs) surrounds the cotyledon. During cotyledon growth, PCs undergo a unique multi-polar expansion going from near cuboidal cells towards interdigitated puzzle-shape cells ^1^. The PC interdigitation event occurs with specific auxin signaling coordination throughout the cotyledon ^2^. The small molecule phytohormone auxin acts as both a local and global signal to coordinate the PC interdigitation ^3–6^. Local auxin activates cell surface signaling cascades that are independent of auxin-induced gene transcription and directly leads to PC interdigitation by regulating ROP GTPase-dependent cytoskeletal reorganization ^1,3,4,6,7^. Auxin maxima and gradients peaking in a specific region of developing organs are a common feature in plants ^8,9^.It serves as guidance for proper patterning, growth regulation, cell type differentiation and organogenesis ^10^. Auxin accumulation at the cotyledon’s tip is a widely reported observation ^11^, however, the mechanism for the generation of this tip-high auxin remains unknown.

At the base of auxin maximum/gradient formation is the integrated regulation of various mechanisms impacting cellular auxin levels ^12,13^, including PIN-mediated polar auxin transport ^14–16^, AUX1-mediated auxin influx ^17^, ABC transporter mediated auxin efflux ^18,19^, auxin biosynthesis and metabolism ^20–22^ and local subcellular sequestration of auxin or its precursors ^23–25^. The absence of auxin transport and/or auxin biosynthesis can partially or completely disrupt the auxin gradients or maxima, proper patterning, and development. Thus, important developmental information is engraved in the spatial distribution of auxin ^8,26–28^, but the nature of this information and the mechanism by which it controls the formation of specific auxin gradients/maxima are still enigmatic.

Auxin is synthesized predominantly from tryptophan through the intermediate INDOLE 3-PYRUVIC ACID (IPyA) in the TAA/YUCCA-regulated IPyA pathway ^29^. Additionally, the INDOLE BUTYRIC ACID (IBA) pathway can locally boost auxin levels through the metabolic conversion of this auxin precursor into active auxin INDOLE ACETIC ACID (IAA). The IBA-to-IAA conversion pathway has been implicated in several developmental processes such as cotyledon expansion, organ size determination and lateral root formation ^25^. It occurs through several enzymatic reactions in the peroxisome where the function of enoyl CoA-hydratase 2, ECH2, is a limiting step ^21^. Cotyledons of the *ech2ibr1,3,10* quadruple mutant with impeded IBA-derived auxin exhibit severely impaired PC morphogenesis ^2,30^. Cytoplasmic IBA levels can also be locally reduced by TOB1 (TRANSPORT OF IBA 1)-mediated vacuole transport ^24^. Subcellular compartmentalization of IBA has been shown to regulate morphogenetic processes such as lateral root formation ^24^.

Despite our extensive knowledge on the critical contribution of both auxin biosynthesis and polar transport to the formation of auxin maxima, it is unclear how these processes are regulated and coordinated to generate and maintain local accumulation of auxin. Here we address this question by using the unique cotyledon system, in which the auxin maximum at the cotyledon’s tip is thought to guide its expansion. We showed that at the cotyledon edge, a file of elongated marginal cells accumulates polarly localized PIN2 forming a polar auxin transport system towards the tip. This auxin transport system is only transiently established via a self-activating and self-terminating mechanism to robustly coordinate PC interdigitation.

## Results

### PIN2-expressing margin cells coordinate pavement cell interdigitation

In developing organs, auxin maxima are formed via the polarized action of the plasma membrane (PM)-localized PIN-family auxin transporters ^8,15^. Published transcriptional data reveals that PINs are induced during germination. Among *PINs*, *PIN2* shows the strongest expression in early imbibed seeds. (**Extended Data 1 a**). Interestingly, we found PIN2-GFP ^31^ protein accumulated specifically in the lateral borders of early expanding cotyledons, forming a file of contiguous marginal cells (MCs) harboring PIN2 at their apical ends, resembling an “auxin transport pipeline” discharging at the cotyledon tip (**Fig. 1 a** and **Extended Data 2 a**). The strong signal at the cotyledon tip does not reflect true PIN2-GFP but autofluorescence, as confirmed by spectral scanning microscopy (**Extended Data 2 b**). Although we cannot exclude the role of other PINs, the apical localization of PIN2-GFP hints a role for PIN2 transporting auxin toward the tip of cotyledons along their marginal cells.

**Figure 1.**
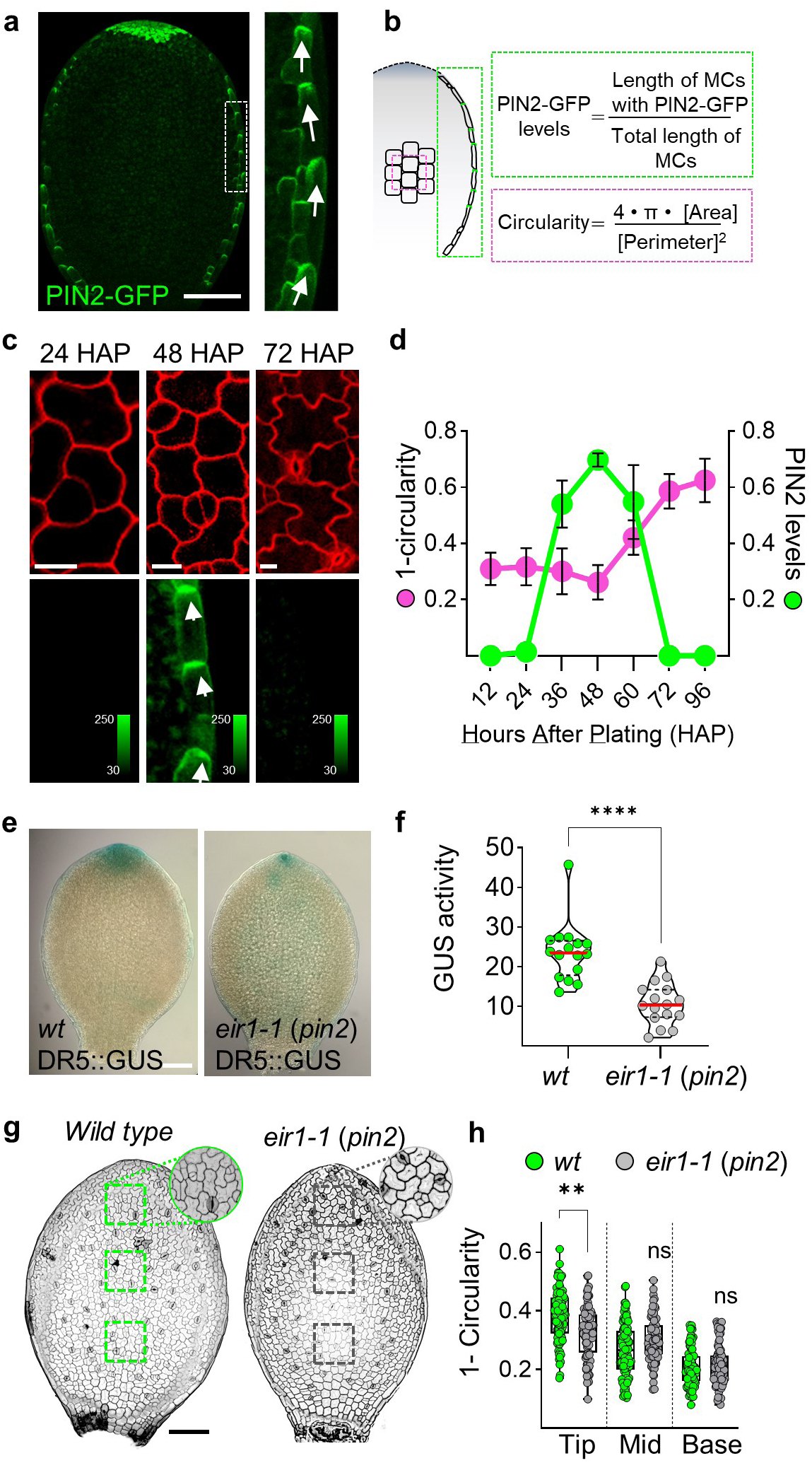
PIN2 accumulation in MCs precedes and plays a role in coordinating PC morphogenesis. **a**, Polar PIN2-GFP in the apical side of marginal cells (MCs) of 48 <u>H</u>ours <u>A</u>fter <u>P</u>lating (HAP) *Arabidopsis* cotyledons. Image corresponds to a maximum intensity projection of a z-stack capturing all perceptible signal. Scale bar = 100 µm. Zoom-in indicates the predicted direction of auxin flow with the white arrows from the base to the tip of the cotyledon. **b**, Schematic of the areas analyzed on each cotyledon. **c-d**, PIN2 accumulates transiently prior to PC morphogenesis. (**c**) Cell shapes visualized by PI (upper panel) staining at the adaxial side of cotyledons expressing PIN2-GFP (lower panel) and grown for 24, 48, 72 hours after <u>p</u>lating (HAP). Shown are representative images captured in the center (PI) and in the border of the cotyledon (PIN2-GFP). Scale bar = 10 µm. (**d**) Quantification of the interdigitation levels (magenta) and PIN2-GFP accumulation level (green) in 12 to 96 HAP cotyledons quantified as indicated in **b.** Interdigitation levels are expressed as 1-circularity, *i.e.*, the greater the value, the greater the degree of interdigitation. Circularity is defined as 4*π* area/perimeter^2^. PIN2 accumulation is quantified as the percentage in length of the border from base to tip that is expressing PIN2. n=12 cotyledons from different seedlings. **e-f**, Reduced auxin transcriptional response in 48 HAP *eir1-1* (*pin2*) mutant. (**e**) *R*educed *<u>DR5::GUS</u>* signal at the cotyledon’s tip in *eir1-1 (pin2)* compared to *wild type*/*wt*. Representative images are shown. The GUS signal was similarly displayed in 95% (19/20) and 84% (21/25) of the cotyledons assayed for *wt* and *eir1-1*, respectively. Scale bar =100 µm. (**f**) Reduced GUS activity in cotyledons *eir1-1* (*pin2*) measured by fluorometric quantification of 4-MU. GUS activity is shown as nanomolar units of 4-MU per sample. *t*-test ***p<0.001, n=18 samples. Sample is defined as the 2 cotyledons in a seedling. **g-h**, Loss-of-function mutant *eir1-1 (pin2)* showed reduced interdigitation in pavement cells at the cotyledon tip. (**g**) Representative images of wild type and *eir1-1 (pin2)* cotyledons. Boxes indicate areas to be quantified. Zoom-in shows the clear reduced interdigitation of pavement cells at the cotyledon’s tip. Scale bar =100 µm (**h**) Interdigitation level (1-circularity) in 3 areas of identical dimensions: tip, middle and base for wild type *Col-0* (green dots) and PIN2 mutant allele *eir1-1 (pin2)* (gray dots) in 60 HAP cotyledons. n=90 cells from 5 different seedlings. *t*-test. ** *p*-value < 0.01

To further understand the correlation between PC interdigitation and PIN2-driven auxin transport, we analyzed the dynamics of PIN2-GFP accumulation in expanding cotyledons, from 12 to 96 <u>H</u>ours <u>A</u>fter <u>P</u>lating (HAP) (**Fig. 1 b**). Strikingly, PIN2-GFP accumulation pattern in cotyledons was transient, only present for a brief period of time (**Fig. 1 c)**. Quantification revealed that PIN2 accumulation starts and consolidates before PC interdigitation is initiated (24-36 HAP) and is depleted once the interdigitation process is completed (72 HAP) (**Fig. 1 d**). This suggests that PIN2-mediated auxin transport along MCs is involved in PC interdigitation presumably through contributing to the cotyledon’s tip auxin maxima.

Importantly, the auxin transcriptional response marker *DR5::GUS* exhibited greatly reduced GUS signal at the cotyledon tip in the *PIN2* loss of function mutant *eir1-1/pin2* ^32^, compared to wild type (**Fig. 1 e**). This reduced GUS signal was corroborated by quantification of GUS activity in a fluorometric assay (**Fig. 1 f**). Additionally, *pin2* showed less interdigitated and smaller PCs at the cotyledon tip (**Fig. 1 g-h and Extended Data 1 c**). Similar phenotypes were also observed in the insertional mutant allele *pin2-T* ^33^ (**Extended Data 1 d-e**). These results imply that PIN2, which is specifically expressed in MCs, has a cell non-autonomous function in the promotion of PC interdigitation in cotyledons. To decipher the functional relevance of MCs in the regulation of PC interdigitation, we laser-ablated all MCs in a cotyledon prior to the initiation of PC interdigitation. After 24 h, cotyledons showed signs of normal physiological status however they displayed a strong inhibition of PC interdigitation (**Extended Data 2 c-g**). The laser ablation experiments hinted a role for PIN2-expressing MCs in the regulation of PC interdigitation. Altogether, 1) low interdigitation in *pin2* cotyledon’s tip, 2) low auxin response in *pin2* cotyledon tips, 3) consistent apical (towards tip) localization of PIN2 in consecutive MCs, and 4) the importance of PIN2-expressing MCs revealed by laser ablation, suggest the importance of PIN2-mediated auxin transport along the cotyledon margins towards the tip for the formation of auxin maxima to coordinate PC interdigitation.

If MCs are indeed transporting auxin, we should be able to detect increased auxin levels in these cells after PIN2 is accumulates. We tested this by comparing nuclear auxin marker before and after PIN2-GFP accumulation (**Fig. 2 a, 24 vs 36 HAP**) in stable non-segregant lines containing both PIN2-GFP and the ratiometric auxin reporter R2D2 ^34^. R2D2 reports auxin-induced degradation of DII-Venus normalized by an auxin non-responsive version mDII protein (**Fig. 2 b**). The nuclear quantification of R2D2 ratiometric index along MCs showed a ∼50% increment after PIN2-GFP is accumulated (**Fig. 2 c**), indicative of increased auxin levels along MCs. Taken together we conclude that a transient PIN2-based auxin flow through the cotyledons margin contributes to the tip auxin maximum and plays a role in coordinating the PC interdigitation process.

**Figure 2.**
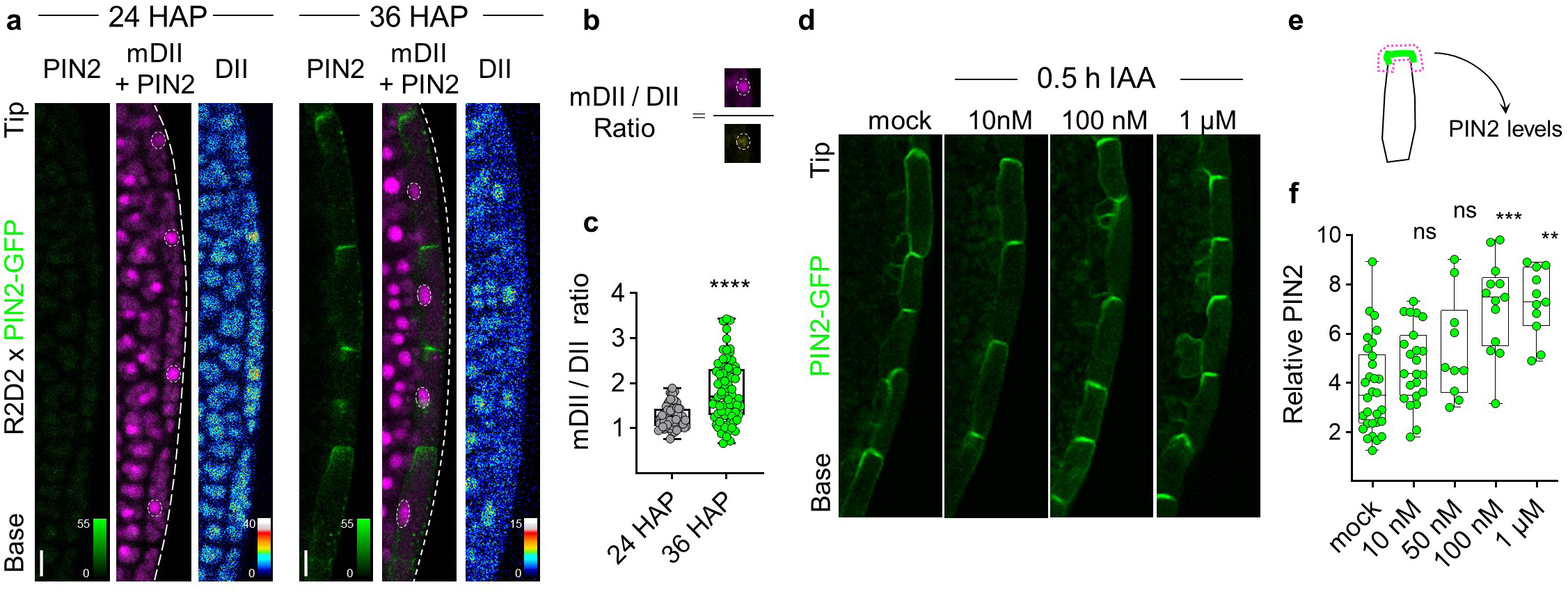
Auxin promotes its own PIN2-mediated efflux along MCs. **a**, Auxin levels increase after PIN2 levels increase. PIN2-GFP+/+ X R2D2+/+ cotyledon’s borders at 24 and 36 hours after plating (HAP). R2D2 signal results from DII-Venus (DII) and mDII-ntTomato (mDII). Scale bar = 10 µm. PIN2-GFP can only be observed on margin cells (MCs) at 36 but not at 24 HAP. **b**, Schematic of quantification of auxin levels using R2D2 in marginal cells (MCs). Nuclear signal intensity from the non-degradable mDII-ntTomato (mDII) is divided by the signal from the auxin-degradable DII-Venus (DII). **c**, Quantification of R2D2 signal according to the formula in **b**. Higher mDII/DII ratio indicate higher auxin response. (n=64 and 75 cells for 24 and 36 HAP, from at least 6 cotyledons, respectively). Box plots indicate 4 quartiles and the median value. *t*-test. **** p-value < 0.0001. **d**, PIN2-GFP-expressing marginal cells (MCs) of 48 HAP cotyledons after 30 min treatment with mock or auxin IAA at the indicated concentrations. **e**, Red dashed lines indicate the shape of regions used for quantification in **f**. **f**, Quantification of apical PIN2-GFP after 30 min of auxin IAA treatment. The mean integrated density obtained in the apical side from MCs is normalized by the background fluorescence. Box plots indicate 4 quartiles and the median value. (n = 10 cell files, each from different cotyledon). *t*-test. *** p-value < 0.001, ** p-value < 0.01.

### Auxin positively feedback regulates polar auxin transport along the MCs

The transient nature of PIN2 accumulation presented an intriguing question: What are the underlying mechanisms behind initiation and termination of the PIN2 accumulation along MCs? The increased auxin levels together with PIN2-GFP polarization is consistent with the well-known effect of auxin regulating PIN expression ^35^. Therefore, we supposed that auxin might positively regulate PIN2-mediated auxin transport along MCs. Indeed, we detected a dose-dependent rapid increase in PIN2 localized to the apical plasma membrane after auxin treatments for 0.5 h, with a significant increase beginning at 100 nM IAA (**Fig. 2 d-f**). PIN2 plasma membrane localization was also more polarized after auxin treatments (**Extended Data 3 d**). Conversely, interruption of auxin transport by NPA treatment disrupted PIN2 polar localization (**Extended Data 3 a-c)** and PC interdigitation (**Extended Data 3 e-f)**. Together these data suggest that an active “tip-wards” auxin transport along the MCs is initially organized by auxin itself.

### IBA-derived auxin regulates PIN2 accumulation along MCs

To understand the biosynthetic origin of the auxin that flows along MCs, we explored the INDOLE 3-PYRUVIC ACID (IPyA) and INDOLE BUTYRIC ACID (IBA) pathways as possible sources of auxin. To this end, we used the presence or absence of key enzymes on each pathway as markers of their respective pathway activity. *ECH2p::YFP-ECH2* ^36^ and cytoplasmic *TAAp::GFP-TAA1* ^11,37^ were used to mark IBA and IPyA pathways, respectively. GFP-TAA1 does not accumulate in MCs prior to PIN2-GFP accumulation (**Extended Data 4 a**), suggesting the IPyA pathway does not initially trigger PIN2 accumulation. Furthermore, transcripts coding for enzymes working in the IPyA pathway (TAA1, TAR2 and YUC1-11) are weakly or not expressed, while those for the IBA pathway (IBR1, IBR3, IBR10, ECH2) are highly expressed at this developmental stage (**Extended Data 4 c-d**). However, GFP-TAA1 did accumulate in MCs and epidermal cells after PIN2-GFP was already accumulated (48 HAP), suggesting that the IPyA pathway may work in reinforcing auxin levels (**Extended Data 4 a, white arrows**). In fact, chemical inhibition of the IPyA pathway reduced the accumulated PIN2 levels by ∼25% but did not inhibit the formation of the PIN2-based ‘pipeline’ along MCs (**Extended Data 4 e-f**).

In contrast, YFP-ECH2 showed clear accumulation in MCs prior to the PIN2 establishment (**Fig. 3 a**) and a clear accumulation at the cotyledon’s tip after PIN2 (**Extended Data 4 b**). ECH2-labeled punctate structures could indicate peroxisomal localization (**Fig. 3 a, white arrows and intensity profiles**), as shown previously in roots ^30^. Peroxisomes is where the IBA-to-IAA conversion takes place ^30^. This punctate localization of ECH2 in MCs suggests the biosynthesis of IBA-derived IAA in those cells. Thus, we anticipated that PIN2 establishment along MCs would be compromised in IBA-to-IAA conversion mutants. Indeed, *ech2ibr10* showed decreased levels of apical PIN2-GFP in cotyledon MCs at 48 HAP (**Fig. 3 b-c**). Consistently with reduced auxin maxima levels, cotyledons of *ech2ibr10* were smaller and hyponastic (**Extended Data 4 g**) and exhibited defects in PC interdigitation, as reported previously ^2,30^ (**Fig. 3 d-e**). Disrupted auxin transport through the cotyledon margins would explain the reduced *DR5::GUS* expression previously observed in the tip of *ech2ibr10* cotyledons, whose severe PC interdigitation defects were rescued by exogenous auxin ^2^. Altogether, these results support the hypothesis that predominantly IBA-to-IAA conversion drives the onset of PIN2 accumulation along MCs, resulting in the formation of the tip auxin maximum that spatiotemporally coordinates PC interdigitation.

**Figure 3.**
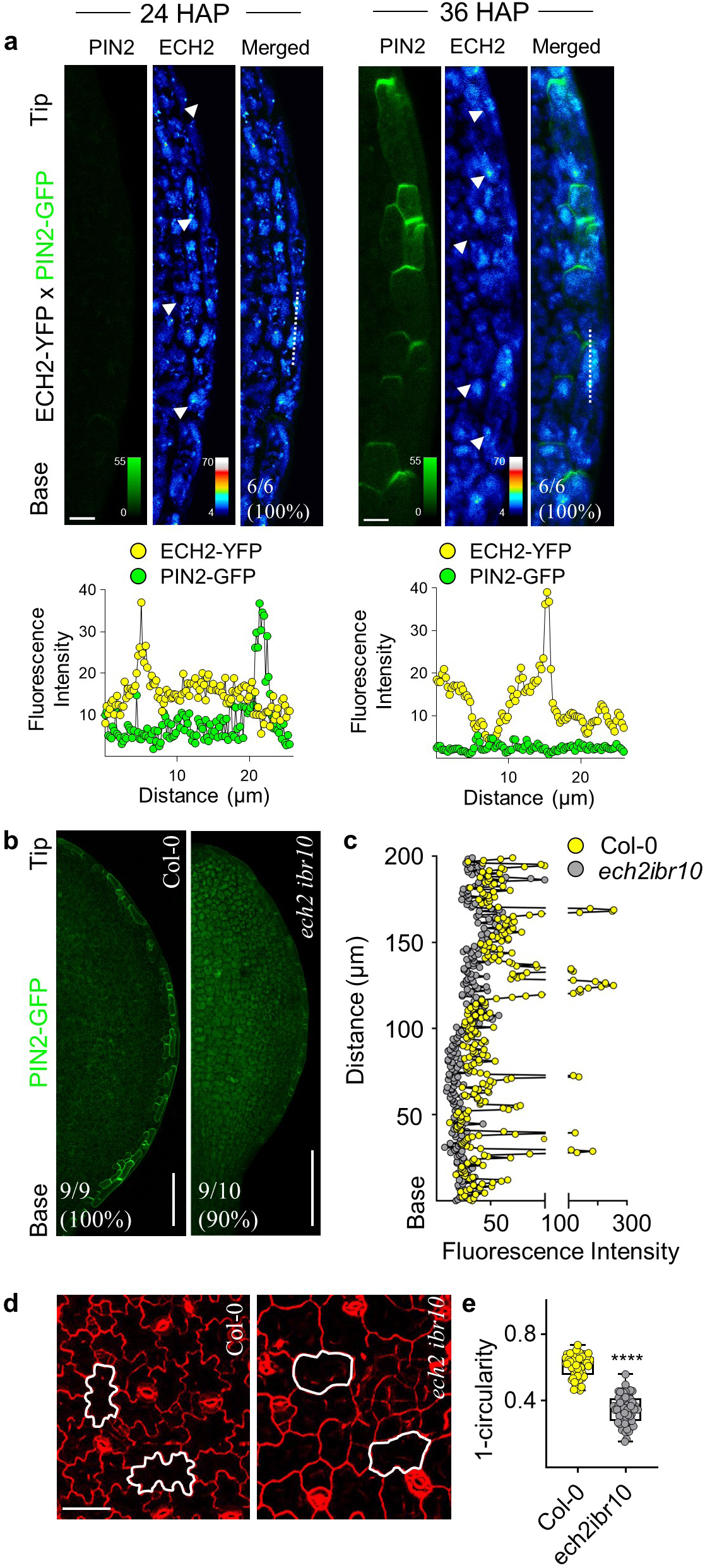
IBA-derived auxin regulates PIN2 accumulation along cotyledons margin cells. **a**, *Upper panel*, ECH2 expression precedes PIN2. ECH2-YFP+/+ X PIN2-GFP+/+ cotyledons were imaged in the absence and presence of PIN2-GFP at 24 at 36 HAP, respectively. YFP-ECH2 (white arrowhead) is present at 24 HAP, before PIN2 accumulation. Phenotype frequency is shown as a percentage of the number of cotyledons observed. Scale bar = 20 µm. *Lower panel*, Quantification of the signal intensity along the dashed line in the upper panel. **b**, IBA-derived auxin is critical for PIN2 accumulation at MCs. PIN2-GFP from cotyledons of wild type *Col-0* and *ech2ibr10 -/-* grown for 48 hours after root protrusion. Phenotype frequency is shown as a percentage of the number of cotyledons observed. Scale bar = 100 µm. **c**, Quantification of PIN2-GFP signal along cotyledons borders of *wt Col-0* (gray) and *ech2ibr10* (yellow) captured from base to tip. **d**, IBA-derived auxin is critical for interdigitation. Pavement cells of 5 days-old cotyledons stained with propidium iodide (PI) to outline the cell borders of wild type *Col-0* and double mutant *ech2ibr10 -/-*. Scale bar = 100 µm. **e**, Quantification of pavement cell shape (Circularity) in wild type and *ech2ibr10 -/-* cotyledons, as shown in **d**. n=60 cells from 6 cotyledons, *t*-test. **** *p*-value < 0.0001.

### The distribution of TOB1 to the tonoplast is associated with PIN2 downregulation

The fact that the apical PIN2 levels were greatly reduced in the *ech2ibr10* IBA-to-IAA conversion mutant prompted us to hypothesize that the PIN2 down-regulation observed around 60-72 HAP in wild type cotyledon MCs (**Fig. 1 d**) may result from a reduction in the IBA-to-IAA conversion in those cells. One possible mechanism for this reduction would be through down-regulation of cytoplasmic IBA availability. Recently, TOB1 was described to reduce the IBA pool by importing IBA into vacuoles, consequently altering lateral root formation ^24^. Thus, TOB1 might also regulate IBA availability in cotyledon’s MCs, considering that TOB1 and PIN2 expression patterns are very similar in shoots, both being expressed in epidermal cells in a highly specific fashion (**Extended Data 5 a**). In fact, the translational reporter TOB1p::YFP-TOB1 (hereafter YFP-TOB1) showed highly specific TOB1 protein accumulation at the cotyledon’s MCs during the first 4 days of seedling growth, with some additional expression in neighboring cells (**Extended Data 5 c**). Notably, the YFP-TOB1 subcellular signal pattern changes during cotyledon growth probably reflecting a change in TOB1 subcellular localization. YFP-TOB1 signal became reticular when PIN2 accumulation peaked (**Fig. 4 a, 48 HAP**), whereas it was mainly vacuolar/prevacuolar (ring-like) when PIN2 disappeared (**Fig. 4 a, 60 HAP**). Importantly, we found YFP-TOB1 co-localized with tonoplast marker protein CFP-Vac-ck marker ^38^ (**Fig. 4 b**) and tonoplast marker dye FM4-64 ^39^ and surrounding the vacuole lumen marker dye BCECF ^40^ (**Extended Data 5 b**), similar to observations in roots ^24^. Therefore, we propose that tonoplastic TOB1 in MCs sequesters IBA into vacuoles and thus functions to terminate PIN2 accumulation to the apical PM of MCs.

**Figure 4.**
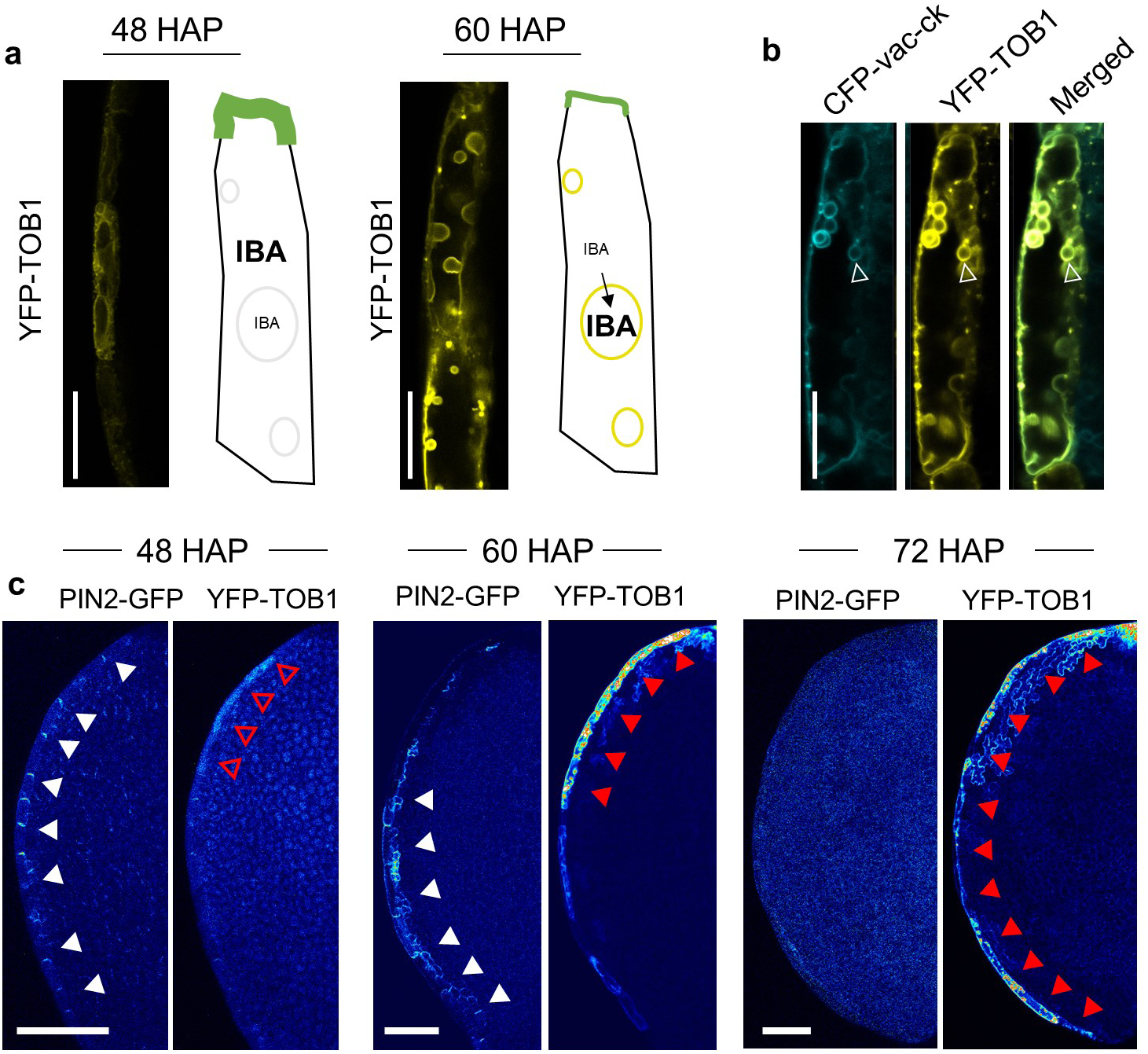
TOB1-mediated vacuole internalization of IBA terminates with PIN2 polar accumulation at cotyledon’s marginal cells. **a**, TOB1 consolidates its tonoplast localization in MCs overtime. TOB1p::YFP-TOB1 (YFP-TOB1) reticular signal at 48 hours after plating (HAP) changes to tonoplast (ring-like) signal at 60 HAP. To the right of each micrograph, a schematic shows the relationship between PIN2 and TOB1 protein levels. At 48 HAP, PIN2-GFP is abundantly expressed whereas TOB1 displays weak reticular expression pattern. At 60 HAP, PIN2-GFP levels are substantially lower and TOB1 shows clear tonoplast accumulation. The complete time course analysis from 24 to 96 HAP can be found in Extended Data 5. Scale bar size = 20 µm. **b**, Tonoplast marker CFP*-vac-ck* signal co-localize with YFP-TOB1 signal in 60 HAP cotyledon’s MCs. White empty arrowhead indicates the superimposition of CFP*-vac-ck* with YFP-TOB1 signal. Scale bar size = 10 µm. **c**, TOB1 and PIN2 accumulation pattern are inversely correlated. Spectral confocal microscopy image of PIN2-GFP +/+ X YFP-TOB1 +/+ cotyledons grown for 48, 60, 72 hours after plating (HAP). White arrowheads indicate the progressive disappearance of PIN2-GFP signal. Empty red arrowheads denote the reticular YFP-TOB1 expression. Red arrowheads indicate the progressive appearance of tonoplast YFP-TOB1 signal. Scale bar size = 100 µm.

### Vacuolar IBA importer TOB1 functions to downregulate PIN2 levels in cotyledons

If TOB1 functions to terminate PIN2 accumulation, then its accumulation is expected to correlate with PIN2 accumulation/disappearance temporal patterns. Thus, we imaged cotyledons expressing translational reporters for both PIN2-GFP (*PIN2p::PIN2-GFP*) and YFP-TOB1 (*TOB1p::YFP-TOB1*) before, during and after PIN2 accumulation in MCs. YFP/GFP signal was accurately separated by means of spectral microscopy ^41^. We found TOB1 signal showed progressive tip-to-base accumulation along the cotyledon’s margin while PIN2 signal disappears in the same direction (**Fig. 4 d, white vs red arrows**). Although we presumed this is part of a continuous process, we noted 3 discrete stages. First, YFP-TOB1 showed weak signal near the cotyledons tip while PIN2-GFP was highly accumulated along the entirety of the MCs file (**Fig. 4 c, 48 HAP**). Second, at 60 HAP, YFP-TOB1 consolidated into a ring-like subcellular signal close to the cotyledon tip, while PIN2-GFP disappeared at the tip but remained at the base (**Fig. 4 c, 60 HAP**). Last, PIN2-GFP was depleted from MCs, whereas YFP-TOB1 was strongly accumulated, opposite to what was found 24h earlier (**Fig. 4 c, 72 HAP**). These results indicate that PIN2 and TOB1 accumulate in MCs in an opposing manner.

Next, we tested if TOB1 is functionally required to downregulate PIN2 by introgressing PIN2-GFP into two loss-of-function alleles, *tob1-1* and *tob1-3* (**Extended Data 7 a)**. Both point (*tob1-1*) and insertional mutation (*tob1-3*) increased cotyledon surface areas (**Extended Data 6 a).** Additionally, both *tob1* mutants showed more expanded epidermal cells when compared to equivalent wild-type cotyledons (**Extended Data 6 b-d**). As expected, absence of TOB1 causes a 25-30% increase in PIN2 levels in 60 HAP cotyledons (**Fig. 5 a**) and also a delayed PIN2 downregulation observed at 72 HAP (**Extended Data 6 e**). To test the biological significance of this increment on PIN2 levels, we evaluated the auxin response distribution pattern as reflected with *DR5::GUS* expression. In both *tob1* alleles, *DR5::GUS* signal was enhanced at the tip and at the edges compared to *Col-0* wild type (**Fig. 5 c**). Increased *DR5::GUS* signal in *tob1* was indicative of increased upwards auxin transport and reflecting the inability to prevent IBA to IAA conversion by IBA vacuole storage (red dashed lines and arrowheads in **Fig. 5 c**). Interestingly, in root tips the PIN2-GFP levels remain unaltered in *tob1-1* and *tob1-3* mutants (**Extended Data 7 b-c**). Additionally, we did not find evidence for genetic interaction between TOB1 and PIN2 for controlling root gravitropism, as *tob1-1 eir1-1* double mutant shows a similar agravitropic response as *eir1-1* mutant (**Extended Data 7 d-e**). Consistent with these findings, we found that TOB1 and PIN2 are not accumulated in the same cell type in roots (**Extended Data 7f**), further corroborating the notion that the TOB1 regulation of PIN2 levels is a cell-autonomous phenomenon. Thus, our results showed that in cotyledons but not in roots TOB1 functions to downregulate PIN2 levels most likely by sequestering IBA into vacuoles and consequently reducing cytoplasmic IAA levels.

**Figure 5.**
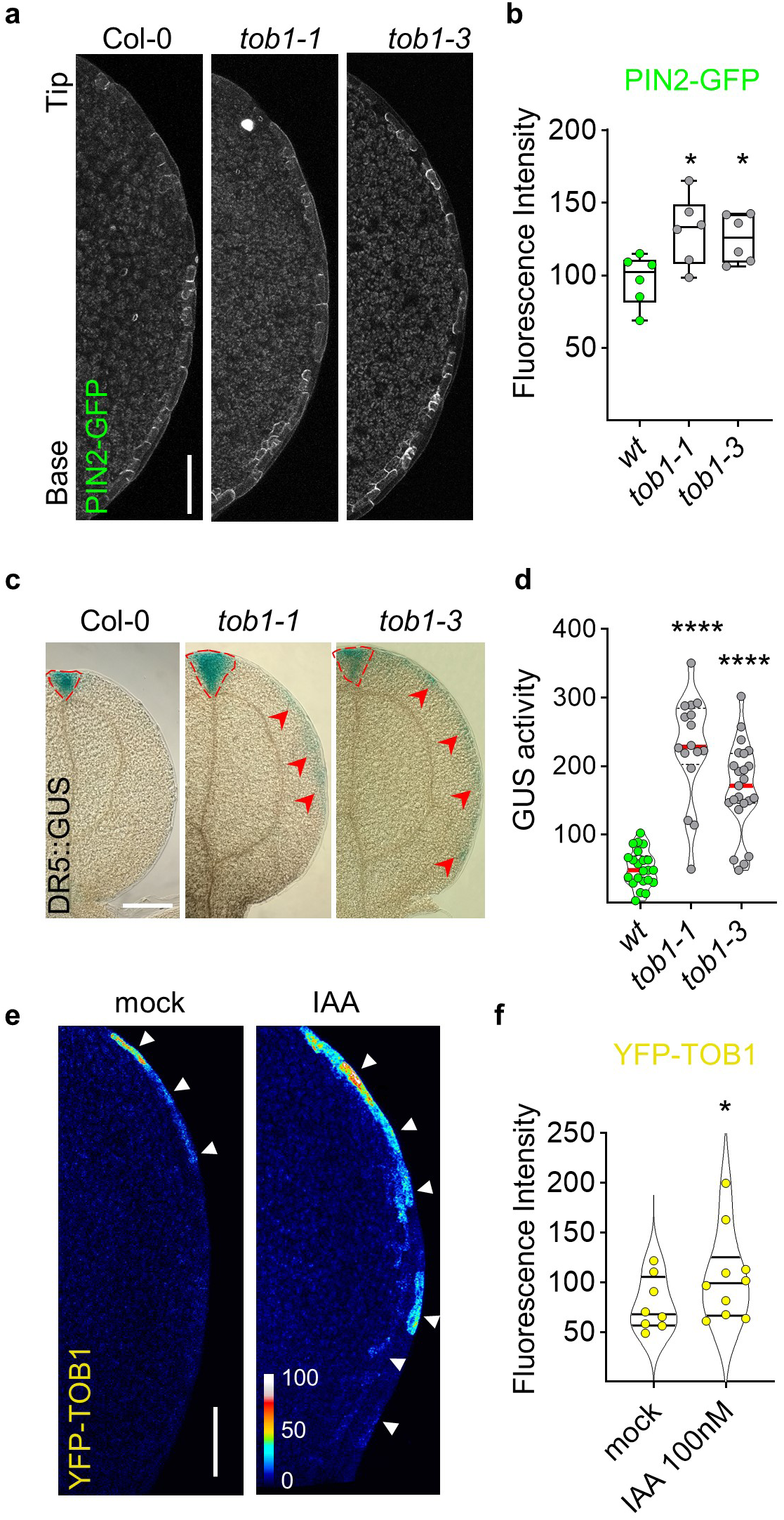
TOB1 regulates PIN2 levels in Arabidopsis cotyledon marginal cells. **a**, PIN2 levels increase in *tob1* mutants. Confocal images of PIN2-GFP cotyledon in *Col-0, tob1-1-/-* and *tob1-3-/-* mutant background, generated by introgression. All lines are non-segregant F3 PIN2-GFP+/+. Images are z-stack maximum projection capturing all detectable fluorescence signal. Scale Bar = 100 µm. **b**, Quantification reflecting PIN2-GFP levels in *tob1* mutants. For quantification, a region of interest was defined around the entire file of MCs. Box plot graph shows data dispersion and the median value. *t*-test, * p-value < 0.05, (n=6 from different 60 HAP cotyledons). **c**, Auxin transcriptional response distribution pattern revealed by DR5-GUS histochemical analysis in 96 HAP cotyledons of wild type, *tob1-1* -/- and *tob1-3* -/-. All lines are non-segregant F3 DR5-GUS +/+. Scale bar = 400 µm. **d**, GUS activity in cotyledons *tob1-1* -/- and *tob1-3* -/- measured by fluorometric quantification of 4-MU. GUS activity is shown as nanomolar units of 4-MU per sample. *t*-test ***p<0.0001. n=16-23 samples. Sample is defined as the 2 cotyledons in a seedling. **e**, Auxin induced TOB1 accumulation. TOB1p::YFP-TOB1 cotyledons grown for 24 HAP were treated during 4 h with mock or 100 nM IAA. Scale bar = 100 µm. **f**, Quantification of fluorescence intensity of YFP-TOB1 along the cotyledon’s MCs upon auxin treatment. Images are z-stack maximum projection capturing all detectable fluorescence signal *t*-test * p < 0.05, n=9 cotyledons.

### Auxin feedback regulates TOB1 expression

The accumulation of TOB1 in the tonoplast (60 HAP) occurs after PIN2-mediated auxin flow throughout the MCs has started (24-36 HAP). Thus, we hypothesized that once auxin accumulates to relatively high levels it regulates TOB1 expression to terminate PIN2-mediated auxin transport. To test this, we treated cotyledons with incipient TOB1 levels (at 24 HAP) with 100 nM IAA for 4 h. Exogenously applied auxin resulted in a ∼30% increment in TOB1 levels (**Fig. 5 e-f**). Under auxin treatments, YFP-TOB1 levels increased along the entire margin (white arrowhead **Fig. 5 e**) but were much higher in those cells closer to the tip (red arrows). This finding is consistent with the above-described tip-to-base wave in the expression of YFP-TOB1 (**Fig. 4 d, red arrowheads**). *In silico* analysis of the *TOB1* promoter revealed several auxin-related motifs, listing ARF7, ARF8 and mARF34 (ARF7/ARF19 paralog in maize) with high confidence in two different algorithms (**Extended Data 7 a**). These observations suggest auxin may induce *TOB1* expression directly, in addition to cytokinin, a well-known *TOB1* regulator ^24^. Alternatively, auxin may regulate *TOB1* gene expression indirectly through auxin-cytokinin interaction. Altogether, we propose that PIN2-transported auxin self-terminates PIN2 expression and auxin transport toward the cotyledon tip by activating the accumulation of TOB1, which further sequesters IBA into vacuoles.

## Discussion

In general, the developmental program of an organism encompasses a continuum of discrete phases, and therefore relies on transient cell-surface signals, transient intracellular signaling structures ^42,43^ and transient cellular arrangements ^44^. Here we show that a transient self-organizing PIN2-based auxin flow takes place in the cotyledon’s margin cells, contributing to the cotyledon’s tip auxin maxima and the PC interdigitation., In leaf, PIN1, PIN3, PIN4, and PIN7 function in vasculature development and independently from PIN2 ^45^. Future research needs to demonstrate if PIN2 establishes tip auxin maxima independently or collaboratively with other PINs.

Self-organizing transient signals, similar to the one described here, may be instrumental during any developmental transition in plants and animals. Beside leaf development, auxin-related self-organizing transient signals may be involved in phyllotaxis ^46^, root elongation, and root branching by regulating the cell proliferation to differentiation transition^47^, or in the transition to rapid hypocotyl elongation during germination ^48^. In animals, self-organized transient signals might control the circumferential actin filament bundles during the embryo to elongated worm transition in *C. elegans* ^49^ or the formation of the multicellular rosettes in the epithelial-mesenchymal transition (EMT) during mammalian embryo development ^50^ or mesenchymal invasion during cancer metastasis ^51^.

Remarkably, this transient auxin flow is generated and dismantled within a 48-hour time frame by positive and negative feedback, respectively. First, the transient auxin flow is self-promoted by turning-on PIN2 accumulation through the biosynthesis of IBA-derived auxin. Subsequently, the local high auxin at the tip generated by the auxin flow induces TOB1 expression, which import IBA into vacuoles, terminating PIN2 accumulation (**Fig. 6**). This self-organizing mechanism secures a pulse of auxin flow towards the cotyledon tip, and thus our findings uncover an exciting mechanism for the generation of transient developmental signals.

**Figure 6.**
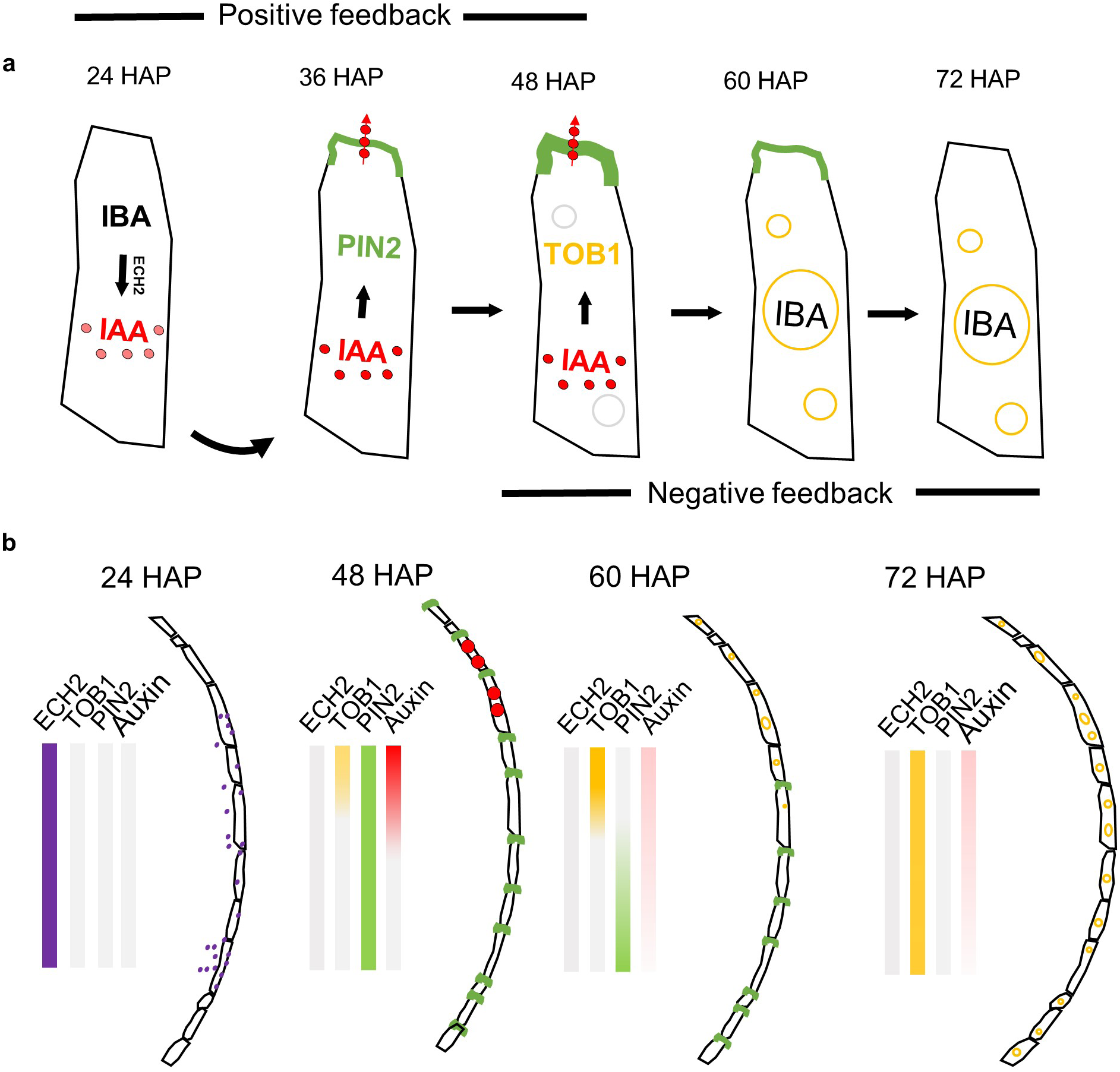
A working model for self-organizing transient auxin flow at cotyledon’s margins. Cotyledon’s margins display a transient self-organized PIN2-mediated auxin transport towards the tip. **a**, Schematic at a cell level. First, a positive feedback (24-48 HAP) via IBA-derived auxin promotes PIN2 expression and PIN2 apical polar accumulation Then, a negative feedback (60-72 HAP) is turned on where auxin induces the expression of TOB1 to sequester IBA into vacuoles, reducing IBA-derived auxin and leading to down regulation of PIN2 and cessation of the auxin flow. **b**, Schematic at a tissue level. At 24 HAP, ECH2 is accumulated along the margin (purple) before PIN2 (green), catalyzing IBA-derived auxin (red). PIN2-based auxin transport moves auxin towards the cotyledon tip. At 48 HAP, higher auxin levels at MCs closer to the tip (red gradient upwards) triggers expression of TOB1 (upwards yellow gradient). TOB1 localizes to the tonoplast gradually from tip-to-base. Tonoplast TOB1 reduce IBA-derived auxin, leading to the reduction of PIN2 levels (downwards green gradient). At 72 HAP, the auxin transport system is completely shut down (gray arrow), which coincides with strong tonoplast TOB1 (dark yellow). At this point, a fully interdigitated field of pavement cells is observed.

Our findings also raise some interesting questions about auxin synthesis, auxin flows, and signaling during developmental transitions. First, what is the advantage of employing IBA over other IAA sources? We showed the IBA-to-IAA pathway has a predominant role in MCs and gene expression indicates a preference for this pathway in the entire matured embryo. Early development and expansion of the matured embryo is fueled through fatty acid β-oxidation, a process similar to IBA-to-IAA conversion ^52^. Thus, the pathway for auxin synthesis may be tightly linked to the seedling energy sources. Future studies could be directed at determining how or if auxin synthesis pathway preferences switch once seedlings become autotrophic. Second, what is the initial signal that activates the IBA-dependent auxin biosynthesis? The signal must be intrinsic as it is produced during seed imbibition. It could be a mechanical signal resulting from increasing turgor pressure and/or initial cotyledon expansion. Third, what is the signal establishing apical PIN2 polarity along MCs?. Different options can be considered; a pre-established tissue-wide polarity field ^53,54^, a combination of gene expression waves ^55,56^ and positive auxin feedback ^57,58^, or a nested self-organizing system between localization and auxin flow, as subjected by PIN1 during PC morphogenesis ^4,59,60^. In fact, PINs are transiently auxin-induced ^61^ and present during leaf serration ^54,62^.

Leaf borders, in particular leaf serration, have been found critical for the evolution of leaf shape diversity ^54,62^. As we have observed polar PIN2 in cotyledons, it would be interesting to decipher if the PIN2-based auxin transient flow and its contribution to PC morphogenesis could work as a link between PC shape diversity ^63^ and leaf shape diversity ^64,65^.

Finally, it is well known that roots and stems show coordinated localization of polar PINs but usually in a multi cell-type context and without a clear start of the process. In contrast, cotyledon PIN2 localization occurs in only one cell type, the marginal cells, and the whole process is activated specifically at 24 HAP, moving auxin tip-wards throughout 10-15 cells. Thus, we propose that transient auxin flow at MCs might bring an exceptional opportunity to study the mechanisms underlying PIN-based auxin flow.

## Methods

### Plant Growth Conditions

All plants were grown at 22°C under a 16 h light/ 8 h dark photoperiod with 85-90 µmol/s^-2^ of light intensity. *In vitro* growth was carried out in 0.5X MS basal salt medium with vitamins, 1% sucrose, pH 5.7 - 5.8 and 0.8% Agar (Sigma-Aldrich® A1296). <u>H</u>ours <u>A</u>fter <u>P</u>lating (HAP) are counted from the moment the seeds are transferred to the growth conditions after stratification.

### Plant Materials

Arabidopsis Columbia (*Col-0*) ecotype is the genetic background for the subsequent lines, including mutants *eir1-1/pin2/*CS8058 ^32^, *pin2-T*/SALK_122916C ^33^, *tob1-3/*SALK_205450 ^24^, *tob1-1* ^24^, as well as the reporter lines TOB1p::YFP-TOB1 ^24^, PIN2p::PIN2-GFP ^31^, TAA1p::GFP-TAA1 ^37^, 35S::GFP-LTI6b^66^, and CFP-*Vac-ck*/CS16256 ^38^. R2D2 is in the Columbia-Utrecht ecotype ^34^. Dual reporters and reporters in mutant backgrounds were generated by introgression. Upon crossing reporters with a mutant background, homozygous mutations were confirmed in F2, and non-segregating lines for the reporter were obtained by screening several F3 lines. Only non-segregant F3 lines were used throughout the study. Primer sequences and genotyping strategy can be found in Supplementary Table 1.

### Seed sterilization

For experiments analyzing specific developmental time points, seeds were bleach-sterilized as follows: 10 min agitation with 30% bleach in water with 0.1% TX-100 (Triton X-100), followed by rinsing 5 times with sterile water and subsequent stratification for 48 h at 4°C in dark. Seed sterilization is crucial for the reproducibility of the developmental timepoints described in this manuscript. For routine experiments, sterilization was performed overnight using the vapor-phase method (Clough and Bent, 1998) carried out by simply placing seeds in a desiccator next to a beaker containing 80 mL of bleach, with 1 mL of concentrated HCl carefully added into the bleach ^67^.

### Developmental stages

Using the bleach sterilization protocol and our described plant growing conditions, we observed the following hallmarks in seedling growth: at 0 HAP, a swollen seed coat; at 24 HAP, a longitudinal rupture in the seed coat; at 36 HAP, radicle protrusion but cotyledons remain inside the seed coat; at 48 HAP, cotyledons were greenish and some were covered with their seed coat and root hair-like structures became noticeable at the root-hypocotyl junction; at 60 HAP, green ellipsoidal cotyledons were rarely covered by the seed coat and form a ∼90° angle between them; at 72 HAP, green globular-shaped cotyledons were flat with a ∼180° angle between them. *<u>H</u>ours <u>A</u>fter <u>P</u>lating* (HAP) time is counted from the moment seeds are placed in growing conditions after stratification. Developmental time in *ech2ibr10* mutants was timed as hours after root protrusion, as these mutants showed intrinsic delayed germination.

### Vector construction

The *ECH2* upstream regulatory region was amplified using promECH2-EcoRI, (GAATTCGGTTAGATATAGAAATGACGG) and promECH2-PmlI (CACGTGCCGATCAGGATTAGAGCTCAGC) primers. The resulting 577-bp product, spanning the region between ECH2 and its most upstream neighboring gene, was then cloned into the pCR4 vector (Life Technologies) to create pCR4-ECH2prom. The *ECH2* upstream regulatory region was excised from pCR4-ECH2prom using EcoRI and PmlI enzymes and subcloned into the pMCS:YFP-GW ^68^ vector to create pECH2:YFP-GW. The ECH2 cDNA was recombined from pENTR-ECH2 ^30^ into the pECH2:YFP-GW vector using LR Clonase (Life Technologies) to form pECH2::YFP-ECH2, which expresses an N-terminal YFP fusion with ECH2 driven by the *ECH2* upstream regulatory region.

### Imaging and microscopy

Adaxial sides of dissected cotyledons were analyzed. In the early stages of cotyledon development, careful removal of seed coat was necessary to dissect the cotyledon. All images were acquired using laser scanning confocal microscopes (20X objective, NA = 0.75 - 0.8). Specifically, ablation experiments were performed using either a Leica TCS SP5 or a Leica Stellaris 8 microscope. Linear unmixing was conducted using a Leica TCS SP8 X DLS or a Zeiss 880 microscope. Fluorescent protein (FP) signals were captured using the following settings: propidium iodide (PI; excitation 543/561 nm, emission 580-660 nm), GFP (excitation 488 nm, emission 505-535 nm), and YFP (excitation 488/514 nm, emission 492-530 nm or 520-570 nm, respectively). PC shape analysis was performed with PaCeQuant software as described before ^6,69^.

### Linear unmixing

Signal separation of GFP/YFP or GFP/PI (propidium iodide) was performed by linear unmixing of spectral images ^41^ acquired with either the Leica TCS SP8 X DLS or Zeiss 880 Elyra systems, generating identical results. For the Leica TCS SP8 X DLS, signal separation was performed using lambda mode with a 20X/NA 0.75 objective, excitation with a 488nm white pulse laser scanning from 490 nm to 630 nm, and detection with HyD detectors. The images were processed using the linear unmixing algorithm in LASX software. For the Zeiss 880, GFP/YFP and GFP/PI signal separation was performed using spectral imaging mode with a 20X/NA 0.8 objective, MBS 488, excitation with a 488nm argon-ion laser and a 32-channel GaAsP spectral detector capable of simultaneous detection of 8.9 nm bands of signal ranging from 410 nm to 695 nm. For GFP/YFP and GFP/PI separation, we used a detection window of the 16 channels between 490-632. The images were processed using the linear unmixing algorithm in Zeiss ZEN software.

### Image analysis

Intensity measurements and circularity [4π *(area/perimeter^2^)] were obtained using Image J 1.40 (Wayne Rasband, NIH, USA) within a region of interest (ROI) as indicated in each figure legend. Circularity is an isoperimetric quotient, which compares the area of a shape, in our case a pavement cell, against the area of a circle with the same perimeter. Higher circularity values represent less lobing, as they are closer to a circle shape ^70^. To observe direct correlation with lobing, we defined interdigitation as 1-circularity. For cell border detection, dissected embryos were incubated either in 100 µg/mL PI for 10 minutes at room temperature (older cotyledons) or for 10 minutes at 4℃ with 5 µM FM4-64 (younger cotyledons) prior to imaging. For apical PIN2-GFP abundance, we quantified the fluorescence at the apical region and normalized it against the background signal. For PIN2-GFP protein polarity, we quantified the fluorescence intensity at the apical side relative to a region of similar area at the middle of the lateral side for each MC. Similar results were obtained regardless of whether the lateral side was located closer to the tip or the base. The fluorescence intensity was obtained in Fiji as the integrated density from z-stack maximum projection confocal images. Both apical and lateral intensities were normalized against the average intensity of 3 randomly selected ROIs in the background.

### GUS (beta-glucuronidase) activity assays

GUS staining assays using X-gluc as a substrate, also known as histological assays, were performed with slight modifications based on previous reports ^71,72^. In brief, plant samples were fixed in cold 80% acetone for at least 2 h, which could be extended overnight. Subsequently, they were vacuum infiltrated with the GUS solution for 10 min and incubated in the dark at 37°C for 0.5h. The GUS solution was prepared as described previously ^71^. Alternatively, 2-4 h incubation was necessary without vacuum infiltration. After staining, cotyledons were “cleared” in 70% ethanol with agitation overnight to remove the chlorophyll. Finally, cotyledons were dissected and observed under a light microscope.

GUS activity assays in intact plant tissue with 4-MUG as a substrate, also known as fluorometric assay, were performed with adaptations from previous reports ^73,74^. Samples (a pair of cotyledons) were incubated in 96-well microplates (one sample per well) containing 150 μL lysis buffer (50 mM sodium phosphate, pH 7.0, 10 mM EDTA, 0.1% Triton X-100). As suggested in the referenced literature, the entirety of the sample tissue must be completely submerged in the buffer for substrate penetration. Subsequently, 1 mM 4-MUG was added to initiate the reaction, which was then incubated at 37°C for 90 min begore being terminated with 50 μL of 0.2 M Na2CO3. The substrate 4-MUG is converted to the fluorescence product, 4-MU. To measure 4-MU fluorescence intensity, a multi-mode plate reader SpectraMax iD5 was used under the following specific wavelengths; 365 nm for excitation and 455 nm for emission. A standard curve generated with 4-MU (in lysis buffer plus stop solution) corroborated linearity (R2 = 0.99) for this assay under our experimental conditions within the range of 10 nM to 1000 nM of 4-MU. GUS activity is expressed as nanomolar units of 4-MU per sample.

### Chemical treatments

Exogenous treatments with IAA (100 nM, EtOH) were performed by directly diluting them in liquid growth media without phytoagar. To inhibit the IPyA auxin-biosynthetic pathway, a two-chemical cocktail containing 50 µM Yucasin and 10 µM L-Kyneurin was used to ensure maximum inhibitory activity ^75^. Yucasin (10mM, DMSO) and L-Kyneurin (50 mM, DMSO) were directly diluted in growth medium before plating. At 24 HAP, seedlings were transferred and incubated for 12 h. in the presence of the chemicals. For vacuole markers, seedlings at 60 HAP were treated with 5 µM FM4-64 for 10 min and 10 µM BCECF-AM for 1 h in darkness. After incubation samples were washed for 30 sec before mounting for observation. FM4-64 (Stock of 10 mM, DMSO) and BCECF (Stock of 10 mM, DMSO) were directly diluted in liquid growth medium without agar.

### Laser ablation

Seedlings of 35S::GFP-LTI6b ^66^ at 48 HAP were incubated with 100 µg/mL with Propidium Iodide (PI) in a microscope slide. A UV 405 nm pulsed laser (300 µW power) at 100% laser intensity was used for cell ablation in a Leica SP5 confocal system or Leica Stellaris 8. For LeicaSP5, each cell was irradiated for 5 sec in maximum optical zoom. PI penetrating the cell borders and marking the nuclei demarcated successful ablation. The whole file of marginal cells was ablated within 2-3 minutes. For Stellaris 8, a small window at maximum optical zoom was assigned to each MCs and a sequential irradiation was initiated using the multi-acquisition mode. After ablation, seedlings were re-incubated *in vitro* in the same growing conditions for further analysis after 24 hours. Vasculature patterns were observed by mounting the mock and ablated samples in Image-IT™ Plant Tissue Clearing Reagent (Invitrogen, V11328).

### In silico analysis

The sequence spanning from +1346bp until transcription start site (TSS) of the TOB1 gene (AT1G72140) was analyzed using promoter analysis tool PlantRegMap. Additionally, the sequence extending +1000bp upstream the TSS was examined using PSCAN, http://159.149.160.88/pscan, a widely used promoter analysis tool. The PINs and TAA/YUCCA gene expression analysis was built by submitting gene codes to Arabidopsis ePlant browser http://bar.utoronto.ca/eplant. Cell-type specific expression analysis data for PIN2, TOB1 was obtained from translatome data at http://efp.ucr.edu.

### Statistical methods

All statistical analyses were performed using GraphPad Prism®. An unpaired two-samples two-tailed *t*-test was used throughout the study, with a 95% confidence interval: Statistical significance was defined as follows: ns (no significance) = p-value > 0.05, * = p-value < 0.05, ** = p-value < 0.01, *** = p-value < 0.001. Equal variance was assumed for the *t*-test and thus was corroborated with a post F-test.

## Accession numbers

PIN2, AT5G57090; TOB1, AT1G72140

## Author Contributions

P.P-H carried out most of the experiments; Z.Y. and P.P-H interpreted the results and designed experiments. M.M. cloned constructs for TOB1 and ECH2 supervised by L.S. S.N, W.T, X.P, Z.L and C.R. contributed critical skills in microscopy. P.P-H. and Z.Y. wrote the manuscript. All authors commented on the manuscript.

## Acknowledgment

We acknowledge David Carter at UCR microscopy core for technical assistance with Zeiss 880 microscopy, Lei Shi at FAFU-UCR center for technical assistance with Leica SP8 microscopy, and Olivier Brun for technical assistance with Leica Stellaris 8. We are thankful to Dr. Yi Tao for providing published material, TAA1p::GFP-TAA1. This work was in part supported by National Institute of General Medical Sciences to Z.Y. (GM081451).

---

**Extended Data 1.**
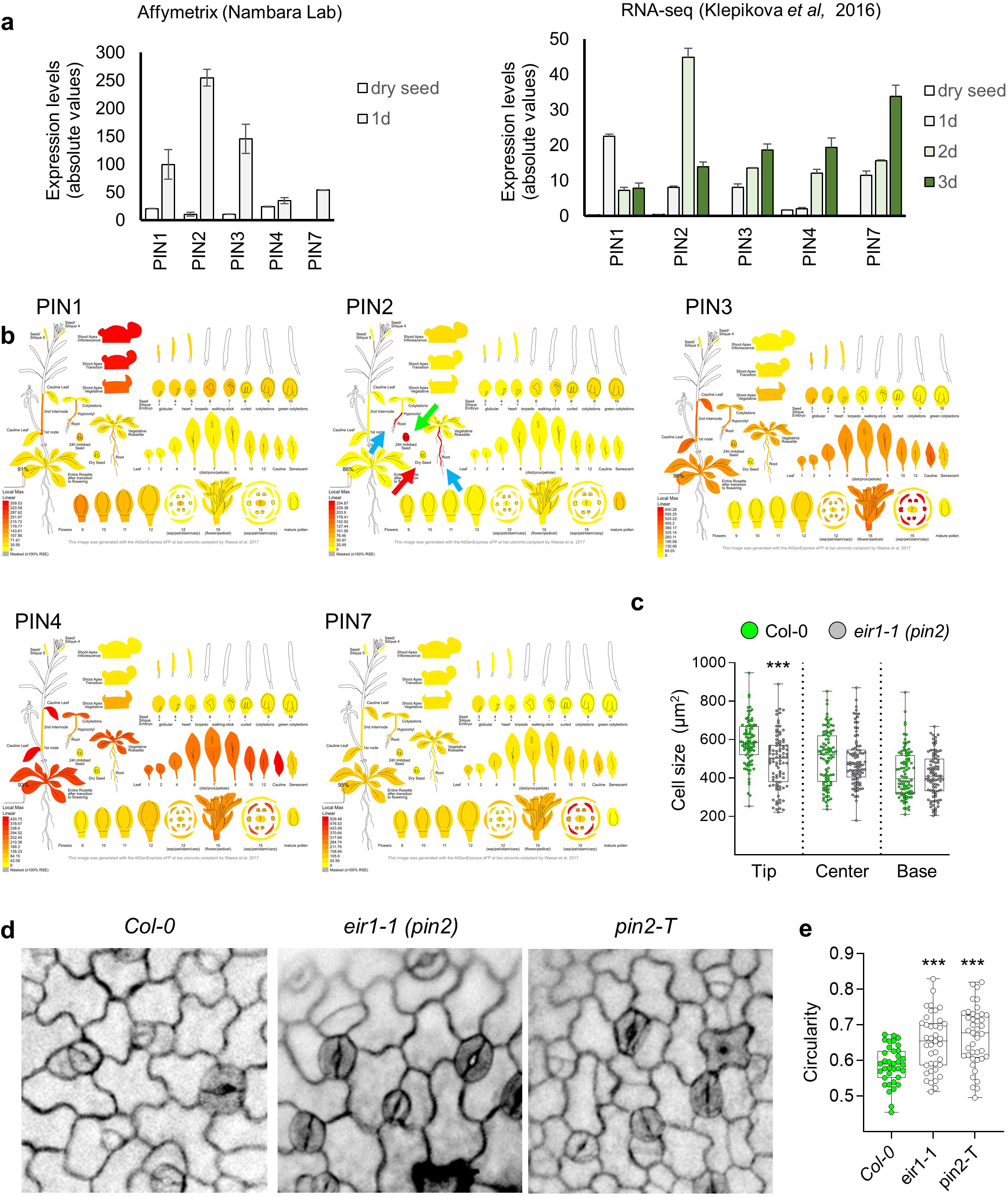
PIN2 is expressed in imbibed seeds and its absence generates reduced PC morphogenesis. **a,** Quantitative expression levels of plasma membrane localized PINs (*PIN1*, *PIN2*, *PIN3*, *PIN4*, *PIN7*) in early developing Arabidopsis; dry seeds (white) and 1, 2, and 3-days old seeds (shades of green). *Left*, Gene expression levels registered by microarray from the Nambara Lab. *Right*, Gene expression levels registered by RNA-seq (Klepikova *et. al.* 2016). Microarray and RNA-seq data are accessible through Arabidopsis *in silico* browser (http://bar.utoronto.ca/eplant). **b,** Developmental map showing transcriptional data for all the PINs localized in plasma membrane; *PIN1/AT1G73590*, *PIN2/AT5G57090*, *PIN3/AT1G70940*, *PIN4/AT2G01420*, *PIN7/AT1G23080* gene codes were subjected data retrieval from Arabidopsis *in silico* browser (http://bar.utoronto.ca/eplant). Notably, *PIN2* is expressed in seedlings and young plant roots (blue arrows) and in seeds 24 h after imbibition (green arrow), but not in dry seeds (red arrows). **c**, Cell size analysis of *eir1-1* epidermal cells showed smaller cells at the cotyledon tip. Cotyledons were grown for 60 HAP and divided for analysis in three regions of interest (ROI) with identical dimensions; tip, center, base. Cell size was obtained for wild type *Col-0* (green bars) and PIN2 mutant allele *eir1-1* (red bars). Cell size is obtained by ImageJ as the area 2D area occupied by a pavement cell. Box plots indicate 4 quartiles and the median value. *t*-test, *** p<0.001. n=90 cells, from different seedlings. This data is complementary to Figure 1h. **d**, PC shape identified at the tip two of loss-of-function mutant alleles for *PIN2*; *eir1-1* (point mutation) and *pin2-T* (insertional mutant, SALK_122916C). **e**, Circularity index of PCs from *eir1-1 (pin2)* and *pin2-T* mutants compared to *wt*. Higher circularity is indicative of lower lobing. Box plots indicate 4 quartiles and the median value. *t*-test, *** p<0.001. n=40 cells, from at least 3 cotyledons each representing different seedlings.

**Extended Data 2.**
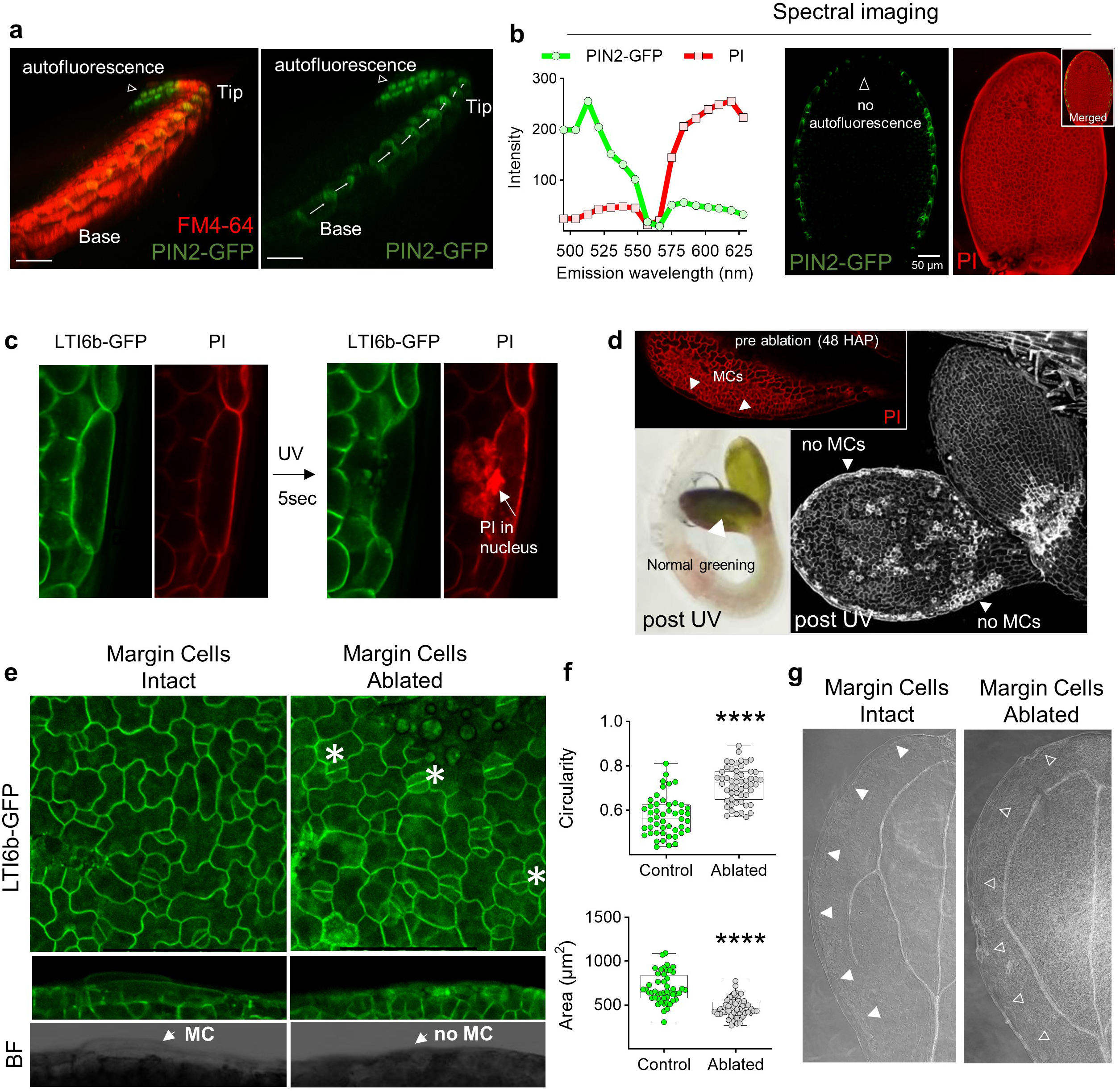
PIN2-expressing margin cells (MCs) play a role during PC interdigitation. **a**, Apical PIN2-GFP at MCs forms an auxin flow “pipeline” towards the tip. PIN2-GFP cotyledons of 36 HAP seedlings were imbibed in 5 μM FM4-64 for 10 min. Z-stacks with 0.3 μm optical slices were used for 3D reconstruction with the Imaris® software. Apical or “tip-facing” PIN2-GFP at MCs can be detected (white arrows) together with autofluorescence at the tip (white empty arrowhead). **b**, Spectral microscopy filters out the autofluorescence at the cotyledon’s tip. PIN2-GFP 48 HAP cotyledons dyed with propidium iodide (PI) were imaged with spectral microscopy collecting 16 channels every 8.9 nm each from 490 to 630 nm. *Left*, GFP and PI emission profiles were used to separate these signals with linear unmixing. *Right,* z-stack projection images displaying isolated PIN2-GFP signal (with no autofluorescence in the tip), isolated PI signal, and an insert with the merged image. **c**, Protocol for laser ablation of MCs. *35S::LTGI6b-GFP* 48 HAP seedlings were mounted on 100 µg/mL vital marker PI. 5 sec of UV laser exposure in the middle of the cell induced PI binding nuclei indicating cell death. **d**, MCs were checked to be alive before UV exposure and dead immediately after UV exposure (pre-ablation). Seedlings turn green after ablation which can be interpreted as no systemic or severe damage after MCs-ablation (post UV – no MCs). **e**, Laser ablation of cotyledon’s MCs severely dampened PC interdigitation. PCs were interdigitated in control cotyledons with fully functional MCs, however, were poorly interdigitated in ablated cotyledons with absent/dead MCs after ablation. The ablation does not affect normal PM-localization of LTI6b-GFP in pavement cells. White asterisks denote normal stomata development in ablated cotyledons. **f**, Quantification of PCs circularity (Q) and cell area (R) from cotyledons shown in **e** (n= 48 and 54 for control and ablated, respectively). *t*-test, **** p<0.0001 **g**, Normal vasculature patterning in cotyledons with ablated MCs. After cell ablation, cotyledons were mounted in Image-IT™ Plant Tissue Clearing Reagent (Invitrogen, V11328) and imaged with darkfield microscopy. Empty arrowhead indica presence of MCs. Empty arrowheads indicate absence of MCs.

**Extended Data 3.**
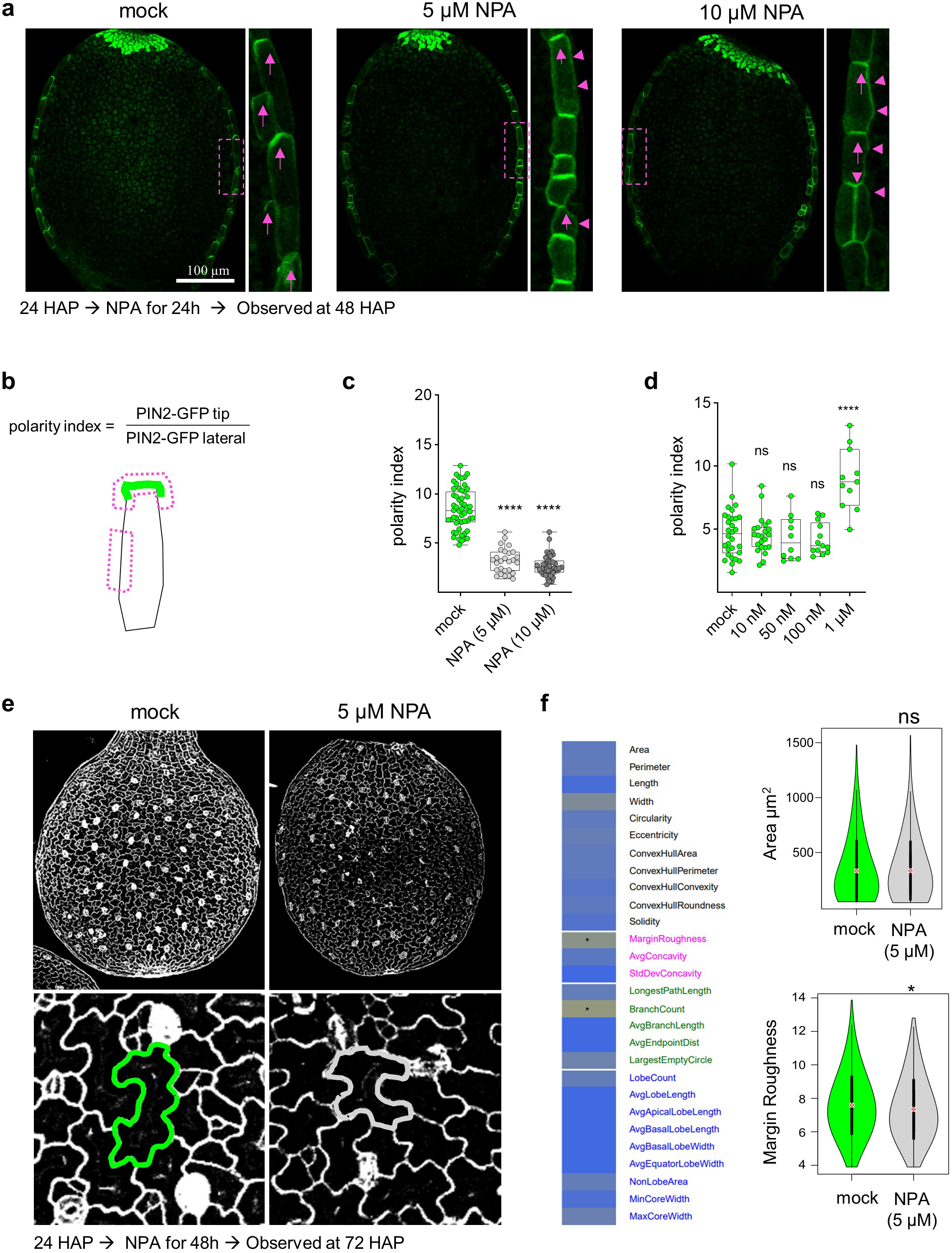
Disruption of auxin transport by NPA treatment reduced both PIN2 polar localization and PC interdigitation. **a,** NPA disrupts PIN2-based auxin flow at MCs. *PIN2p::PIN2-GFP* seedlings at 24 HAP were transferred to 0.5X MS semi-solid media containing mock, (0.01% DMSO), 5 µM or 10 µM NPA and incubated for 24 h before confocal imaging. Zoom-in images in the dash line rectangle are shown. Arrows indicate the normal apical distribution of PIN2-GFP, while arrowheads indicate abnormal non-polar PIN2 distribution caused by NPA. **b**, Schematic representation of the areas considered for the quantification of PIN2 polarity shown in **c**, **d**. **c**, Quantification of the PIN2-GFP polarity index in response to NPA treatments shown in a. n= 60, *t*-test, **** p<0.0001. **d**, Quantification of the PIN2-GFP polarity index in response to IAA treatment for 0.5 h. *t*-test, ns = not significant (*p*-value > 0.05), **** *p*-value < 0.0001. This data complements Figure 2 d-f. **e**, NPA disrupts PC morphogenesis. Wild type seedlings at 24 HAP were transferred to 0.5X MS semi-solid media with mock or 5 µM NPA. Whole cotyledon images (upper panel) and zoom-in images at the center of cotyledons (lower panel) are shown. The superimposed cell contour denotes a much more simpler cell shape observed in NPA treatments. **f,** Shape analysis of PCs in mock and NPA treatments using PaCeQuant software ^69^. *Left*, graphical summary of the NPA effects upon pavement cell shape obtained with PaCeQuant. Yellow indicates reduction in NPA treatments. No asterisk indicates no statistical significance, wilcox-test, * *p*-value <0.05. *Right*, Violin plot the cell area (not significantly different) and Margin Roughness of pavement cells from cotyledons treated with NPA. Wilcox-test, * *p*-value <0.05.

**Extended Data 4.**
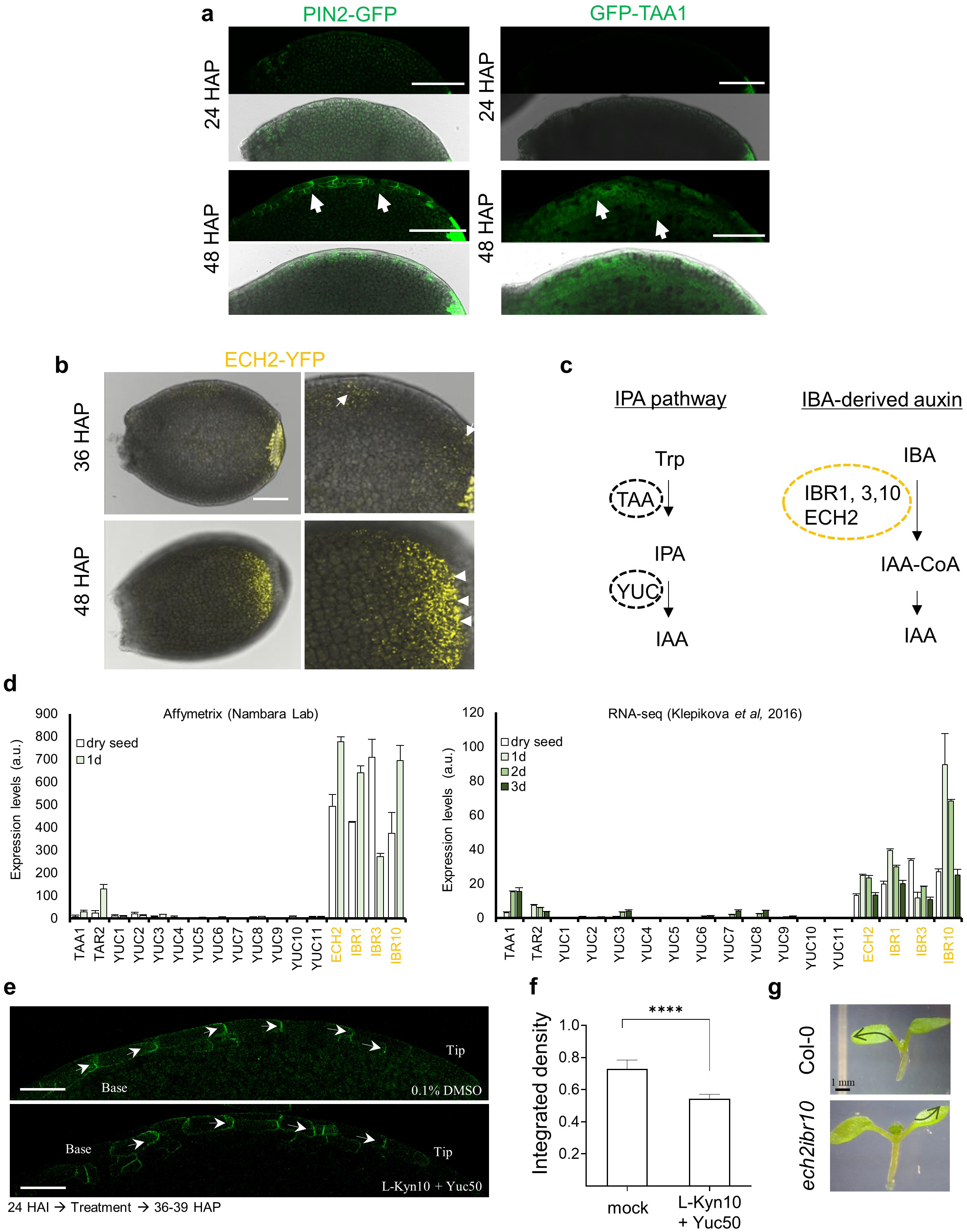
PIN2-mediated polar auxin transport in Arabidopsis cotyledon margin cells (MCs) is initially organized by IBA-derived IAA but not by IAA synthesized via the IPyA pathway. **a**, Border of *PIN2p::PIN2-GFP* (PIN2-GFP) and *TAA1p::GFP-TAA1* (GFP-TAA1) cotyledons grown for 24 and 48 hours after plating (HAP). White arrows indicate PIN2-GFP and GFP-TAA1 signal. Scale bar = 100 µm. **b,** ECH2 accumulation at the tip turns-on after 36 HAP. *ECH2p::ECH2-YFP* cotyledons grown for 36 and 48 HAP merged with bright field channel. White arrows indicate ECH2-YFP signal located at the borders and increasingly at the cotyledon’s tip at 48 HAP. Scale bar = 100 µm. **c**, Schematic of the two main sources for auxin synthesis; the IPyA pathways and the IBA-to-IAA conversion pathway. **d,** Only IBA-to-IAA auxin synthesis pathways genes are abundantly expressed in early developing cotyledons. *Left*, Gene expression levels obtained by microarray from the Nambara Lab at 0 and 1 day (24 h) of growth. *Right*, Gene expression levels obtained by RNA-seq (Klepikova *et. al.* 2016) at 0,1,2,3 days of growth. Microarray and RNA-seq data are accessible through the Arabidopsis *in silico* browser (http://bar.utoronto.ca/eplant). The gene IDs used for this analysis are: TAA1 (At1g70560), TAR2 (At4g24670), YUC1 (At4g32540), YUC2 (At4g13260), YUC3 (At1g04610), YUC4 (At5g11320), YUC5 (At5g43890), YUC6 (At5g25620), YUC7 (At2g33230), YUC8 (At4g28720), YUC9 (At1g04180), YUC10 (At1g48910), YUC11 (At1g21430), ECH2 (At1g76150), IBR1(At4g05530), IBR3 (At3g06810), IBR10 (At4g14430). **e**, Chemical inhibition of IPyA auxin biosynthesis does not disrupt the formation of the polar PIN2 channel. IPyA pathway was chemically inhibited during the initial accumulation of PIN2-GFP at MCs. Seeds of 24 HAP were treated for 12 h in mock or Yucasin 50 µM (Yucca enzymes inhibitor) plus 10 µM L-Kynurenine (TAA enzyme inhibitor). **f**, PIN2-GFP fluorescence intensity around MCs was quantified from treatments shown in e. n=21 cells from 6 cotyledons. *t*-test. **** *p*-value < 0.0001. **g**, Representative shoots from wild type *Col-0* and *ech2ibr10* 8-day old seedlings. Black arrows depict the hyponastic curvature generated in the mutant cotyledons.

**Extended Data 5.**
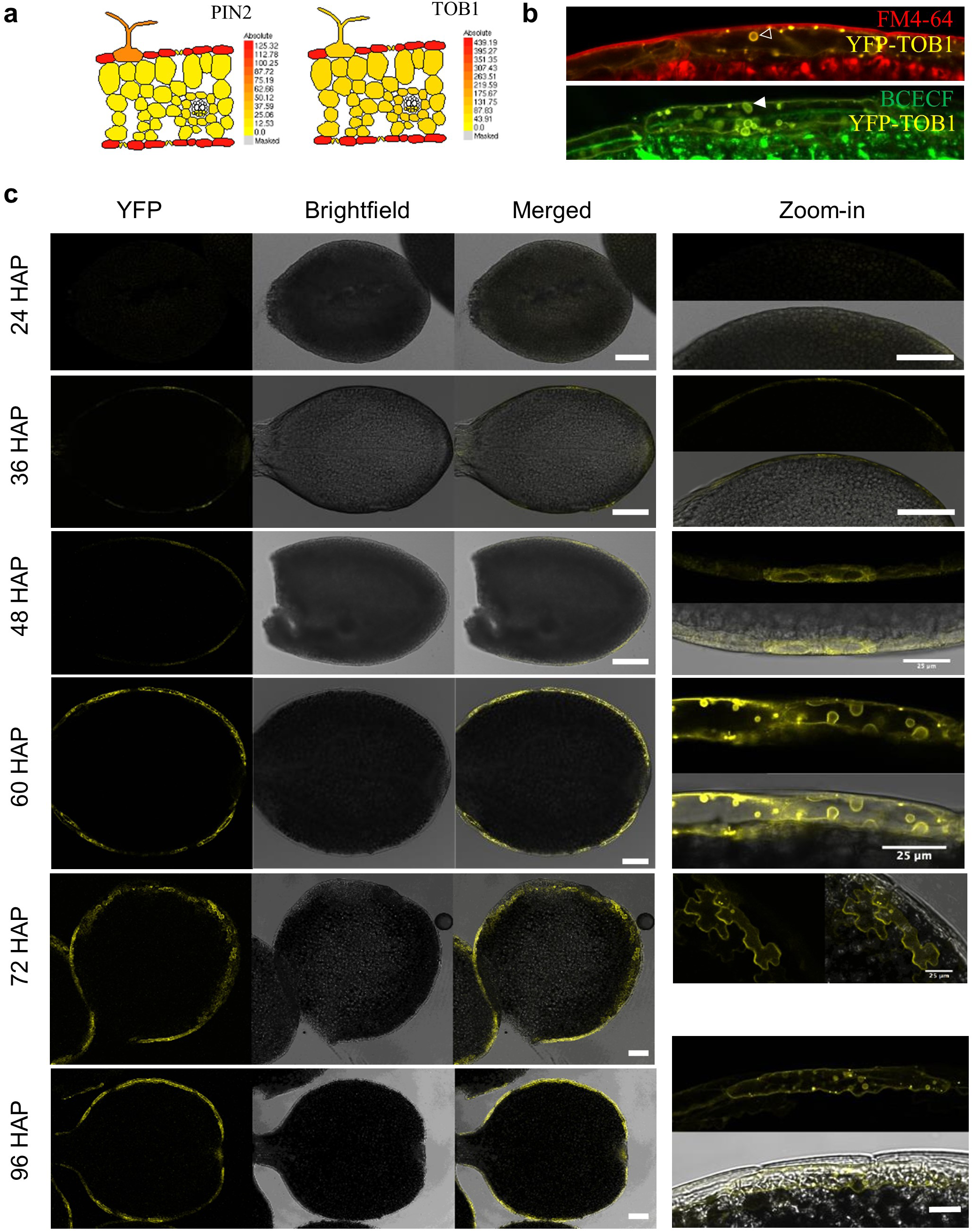
Tissue-wide and sub-cellular localization of TOB1 translational reporter during Arabidopsis cotyledon expansion from 24 to 96 hours after plating (HAP). **a**, Schematic representation of the expression levels in the shoot for IBA transporter *TOB1*/At1g72140) and *PIN2*/At5g57090) (data from http://efp.ucr.edu). *TOB1* expression is high in epidermis (red), emulating *PIN2* expression patterns. **b**, YFP-TOB1 colocalize with the endocytic tracer FM4-64 localized at the tonoplast (white empty arrowhead) and surrounds the vacuole luminal dye BCECF (white arrowhead). **c**, Spatiotemporal analysis of TOB1p::YFP-TOB1 (YFP-TOB1) expression pattern in cotyledons grown during 24, 36, 48, 60, 72 and 96 hours after plating (HAP). The panel displays confocal images (first row), brightfield (second row), merged (third row) and zoom-in images (fourth row). TOB1-YFP displays a clear shift in the subcellular distribution pattern from reticular to tonoplastic (ring-like). Scale bar = 100 µm if it is not indicated except for the zoom-in images in 96 HAP where scale bar = 10 µm.

**Extended Data 6.**
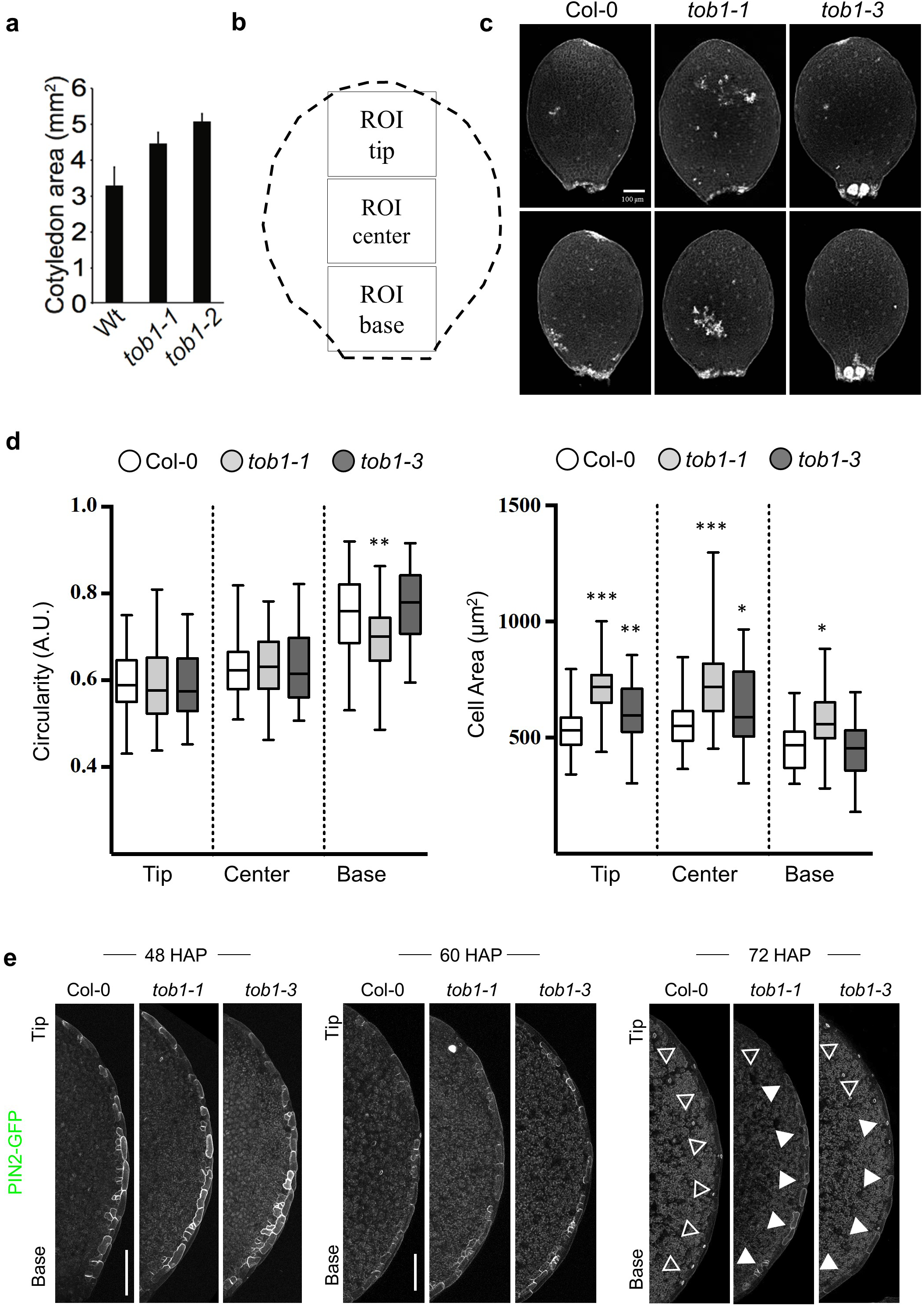
TOB1 regulates cotyledon size also by increasing cell size in early stages. **a,** Cotyledon size of 7 days-old seedlings of *tob1-1* and *tob1-2* mutant alleles ^24^. **b,** Scheme of the display of regions of interest (ROIs) along the cotyledons that are used for quantification. **c**, Representative images of cotyledons from wild type *Col-0, tob1-1 -/-*, and *tob1-3 -/-* using 60 hours after plating (HAP) seedlings. Cell border is detected with Propidium iodide (PI) staining. Cotyledons of similar size were intentionally selected, although notoriously bigger cotyledons were frequently observed among *tob1* mutants. **d**, Quantification of pavement cell circularity and cell size of 60 HAP cotyledons’s tip, center and base. **e**, Spatiotemporal analysis of PIN2 levels in *tob1* mutants. Representative images of PIN2-GFP cotyledon in *Col-0, tob1-1-/-* and *tob1-3-/-* mutant background. Images are z-stack maximum projection capturing all detectable fluorescence signal. Filled arrowhead indicate presence of PIN2-GFP signal. Empty arrowheads indicate absence of PIN2-GFP signal. n =10 cotyledons. Scale Bar = 100 µm.

**Extended Data 7.**
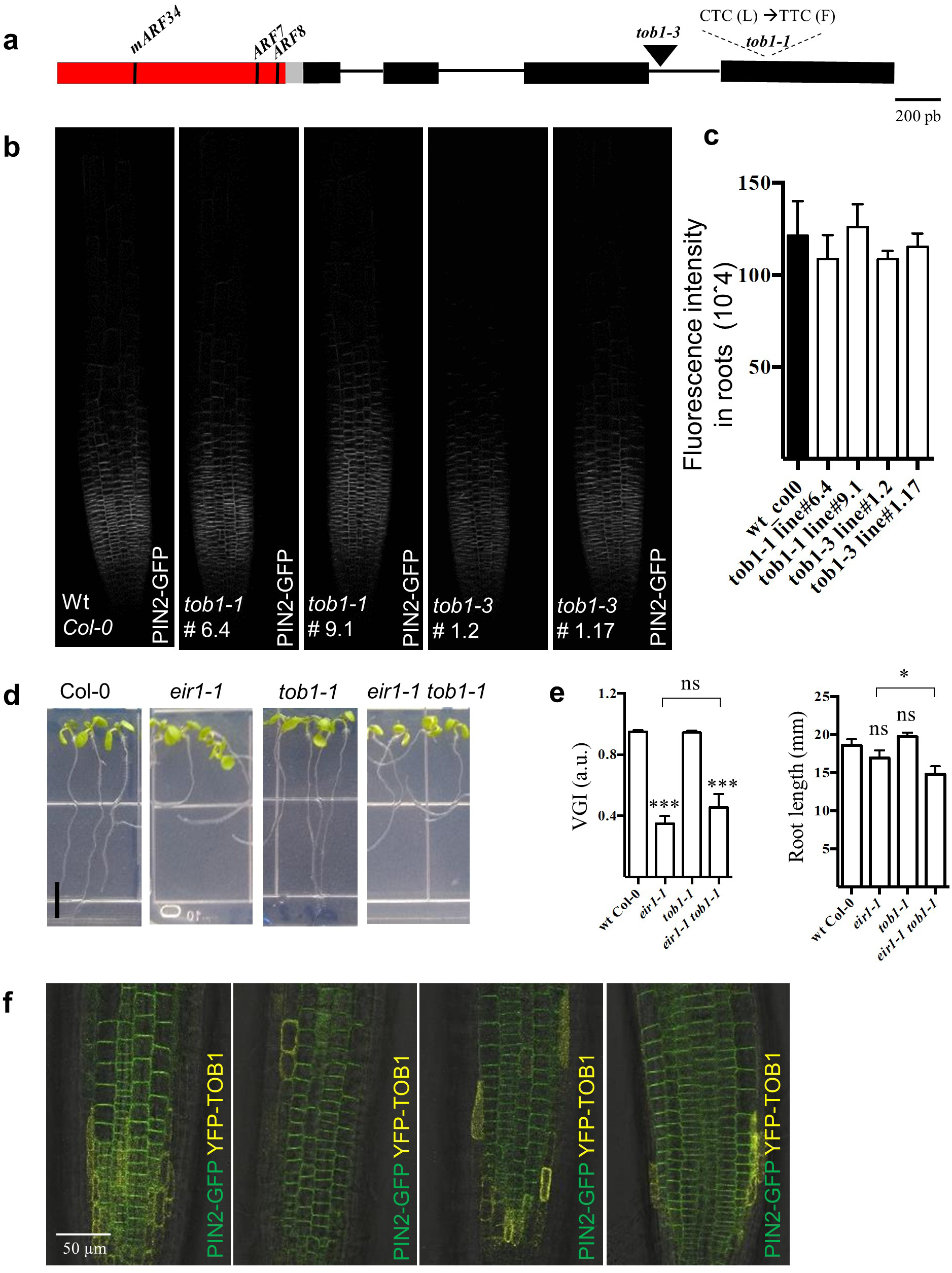
TOB1 does not regulate PIN2 levels in *Arabidopsis* root tips. **a**, Gene model for TOB1 showing t*ob1-1* point mutation (C→T) and *tob1-3* insertional mutation. In the promoter region (red) the relative position for ARF8 binding site is also shown. TOB1 promoter *in silico* analysis revealed putative binding sites are: ARF8 (GCTGTCGCG, position -67, strand -), ARF7 (AGTCACGACGAGAA, position -84, strand -), mARF34 (CTCAGACAACGGT, position -711, strand -). **b**, Confocal images of 3 days old seedling roots displaying PIN2p::PIN2-GFP (PIN2-GFP) in wild type (wt), *tob1-1* or *tob1-3* mutant background. Two different F3 non-segregant lines were used per mutant allele (tob1-x -/- x PIN2-GFP +/+). Images are showing a maximum projection reconstruction from a z-stack. Z-stack captured all focal planes displaying fluorescence signal visible at these settings. **c**, Quantification of PIN2-GFP fluorescence intensity of images shown in b. The whole PIN2-GFP content of the root was quantified using z-stack maximum intensity projections and using the whole image frame as the region of interest of be quantified, n=5 roots. **d**, Images for vertically grown 8 days-old seedlings of wild type *Col-0*, *eir1-1*, *tob1-1, eir1-1 tob1-1* Scale bar = 0.6 cm. Identical results were obtained with *eir1-1 tob1-3* (not shown). **e**, Vertical growth index (VGI) and root length of wild type *Col-0*, the agravitropic mutant *eir1-1 (pin2)*, the point mutant allele *tob1-1* and the double mutant *eir1-1 tob1-1.* VGI quantification was designed to detect weak gravitropic defects ^76^, *t*-test, n =12, ns = no significant, *\* = p*-value < 0.05, *\*\*\* = p*-value < 0.001. Note the notorious decreased in root length in the *eir1-1 tob1-1* double mutant. Statistical comparison is to Wt *Col-0* except when indicated. **f**, Merged-channel confocal images displaying PIN2p::PIN2-GFP (PIN2-GFP) and TOB1p::YFP-TOB1 (YFP-TOB1) signal in the root tip of 4 different 5 days-old seedlings.

