## Supplementary Table 1 for "PIN2-mediated self-organizing transient auxin flow contributes to auxin maxima at the tip of Arabidopsis cotyledons"

Supplementary Table 1. Primer sequence used for genotyping in this study.

| Mutant | Primer name | Primer sequence | Strategy |
|---|---|---|---|
| *eir1-1* | eir1-1_F | 5' TTTGGCCTTGCTTGGTCCCTT 3 | Sequencing with *eir1-1_F* |
| | eir1-1_R | 5' TTACGGCCATCGCAAACCCTG 3' | |
| *pin2-T* | pin2-T_LP | 5' GCAAAAAGGAATCCCAAAATG 3' | PCR with LP/RP and LB1.3/RP pair of primers |
| | pin2-T_RP | 5' TTCAAATGTCCAACGATCCTC 3' | |
| *tob1-1* | tob1-1_F | 5' GCTGTGTTGAGTCTCATTCC 3' | MnlI digestion and sequencing with *tob1-1_F* |
| | tob1-1_R | 5' CTCGACCAAAGCAGCTATTACC 3 | |
| *tob1-3* | tob1-3_LP | 5' CGTGAGTCGTGCGTCTCTCAGG 3' | PCR with LP/RP and LB1.3/RP pair of primers |
| | tob1-3_RP | 5' GATTGATCGTGCGATTGGGACG 3' | |
| *pin2-T and tob1-3* | LB1.3 | 5' ATTTTGCCGATTTCGGAAC 3' | |
